# The rs1421085 variant within *FTO* promotes but not inhibits thermogenesis and is potentially associated with human migration

**DOI:** 10.1101/2021.08.13.456245

**Authors:** Zhiyin Zhang, Na Chen, Ruixin Liu, Nan Yin, Yang He, Danjie Li, Muye Tong, Aibo Gao, Peng Lu, Huabing Li, Dan Zhang, Weiqiong Gu, Jie Hong, Weiqing Wang, Lu Qi, Jiqiu Wang, Guang Ning

**Affiliations:** Department of Endocrine and Metabolic Diseases, Shanghai Institute of Endocrine and Metabolic Diseases, Ruijin Hospital, Shanghai Jiao Tong University School of Medicine, Shanghai, China; Shanghai National Clinical Research Center for metabolic Diseases, Key Laboratory for Endocrine and Metabolic Diseases of the National Health Commission of the PR China, Shanghai National Center for Translational Medicine, Shanghai, China; Shanghai Institute of Immunology, State Key Laboratory of Oncogenes and Related Genes, Shanghai Jiao Tong University School of Medicine, Shanghai, China; Shengjing Hospital of China Medical University, Shenyang, China; Department of Epidemiology, School of Public Health and Tropical Medicine, Tulane University, New Orleans, LA, USA; Department of Nutrition, Harvard T.H. Chan School of Public Health, Boston, MA, USA

**Keywords:** FTO, variant, energy expenditure, obesity, UCP1

## Abstract

Disease-associated GWAS loci are predominantly scattered among noncoding regions of the human genome, which impedes causality estimation. One lead risk signal of obesity–rs1421085 T>C within the *FTO* gene–is reported to functional *in vitro* but lack of organismal evidence. Here, we established global and the brown-adipocyte specific locus-knock-in mice to recapitulate this homologous variant in humans, and discovered the minor allele (C-allele) as one candidate thermogenic locus. Mice carrying the C-alleles showed increased thermogenic capacity and a resistance to high-fat diet-induced adiposity. In terms of mechanism, the knock-in models showed enhanced FTO expression, while FTO knockdown or inhibition effectively eliminated the increased thermogenic ability of brown adipocytes. In humans, the C-allele was associated with lower birthweight, and its allele frequency increases following the environmental temperature decreases. Cumulatively, these findings demonstrated rs1421085 T>C as a functional variant regulating whole-body thermogenesis, and this variation was possibly related to early human migration from hot to cold environments.

## Introduction

Following genome-wide association studies (GWAS), enormous common genetic variants in the human genome have been unraveled and proved to be associated with increasing risk of diseases development (Claussnitzer et al., 2020). However, few of these loci show biological evidence to support their causality with diseases (Claussnitzer et al., 2020; Wijmenga and Zhernakova, 2018). One main challenge is that most positive signals are dispersed within the noncoding regions which are less conserved among different species, hindering further functional studies using genetically modified animal models (Van Der Klaauw and Farooqi, 2015; McMurray et al., 2013) Despite successful applications of gene editing in human cells, functional examinations at the organism level to recapitulate the specific variant in humans are still lacking (Zhu et al., 2019).

To date, the largest obesity-GWAS study has revealed ∼1000 common polymorphisms, explaining ∼6.0% variation of body mass index (BMI) (Yengo et al., 2018). Interestingly, the positive signals repeatedly occur nearby the known monogenic obesity genes, e.g., *MC4R*, *LEPR* and *BDNF* (Van Der Klaauw and Farooqi, 2015). Notably, the strongest signals of the common variants are localized within the introns of the *FTO* gene and show consistent risk-effects across multiple ethnic populations (Locke et al., 2015; Yengo et al., 2018). However, there are no evidence on *FTO* mutations causing obesity in clinical settings (Meyre et al., 2010). Loss-of-function mutations of human *FTO* are lethal owing to serious developmental defects and growth retardation (Fischer et al., 2009). Higher mortality was also observed in global *Fto* knockout mice, and the survived mice exhibited a short stature and a lean phenotype (Fischer et al., 2009). In contrast, the adipocyte-specific *Fto* knockout mice gained more body weight and showed adiposity under HFD compared with WT littermates (Wang et al., 2015). In the past decade, the roles of *FTO* and its genetic variants in adiposity remained a topic of vigorous debate.

A turning point occurred when two research groups independently argued that certain loci (especially for rs1421085) within the *FTO* variant cluster enhanced the transcription of several distal genes (including *IRX3 and IRX5*) but not *FTO per se* (Claussnitzer et al., 2015; Smemo et al., 2014). Very recently, the Nóbrega group reported the functional regulation between three additional functional *FTO* “risk variants” and the candidate genes by using a gene-editing animal model harboring 20 kb deletion of the orthologous obesity-associated interval in the *Fto* gene (Sobreira et al., 2021); while the Claussnitzer and Cox groups further reported a mouse model with a deletion of 82 bp containing the homologous rs1421085 locus in *FTO* which yield a mildly lean phenotype in mice under high-fat diet (HFD) challenge (Laber et al., 2021). Although these genetic editions contain the deletion of the *FTO* rs1421085, they also delete other potential function elements surrounded the *FTO* rs1421085, a precise knock-in mouse model of the *FTO* genetic risk loci (e.g., rs1421085_C-allele) to specifically recapitulate the human variants is still needed to understand the function of the specific risk loci.

Towards this end, we established two global and one brown adipocyte-specific single-nucleotide knock-in mouse models of the homologous human rs1421085 T>C variant, the lead functional variant in the *FTO* gene (Claussnitzer et al., 2015; Peters et al., 2013; Stratigopoulos et al., 2016) (**Figure S1A-D**), and consistently found an enhancement (but not inhibition) of thermogenesis and resistance (but not sensitive) to HFD-induced obesity, at least in part, by increasing FTO expression. Interestingly, the C-allele was associated with lower bodyweight of the newborns, who have functional brown adipose tissue (BAT) as rodents do (Lean et al., 1986). Furthermore, the C-allele frequency was closely related to ambient environmental temperature where multiple ethnic populations settled, with an increasing stepwise pattern from hot Africa shifting to cold Siberia.

## Results

### The homologous rs1421085 T>C variant promotes energy expenditure and resists HFD-induced adiposity

To specifically clarify the biological function of human rs1421085 T>C variant *in vivo*, one global knock-in mouse model carrying a homologous single-nucleotide mutation (the homozygous rs1421085_CC, termed KI^cas9^) was constructed using CRISPR/Cas9 system, with wild-type (WT, rs1421085_TT) littermates as controls (**Figure 1A**, **Figures S1A**-**B**). No significant difference in body weight gain between the two groups was observed when fed a normal chow-diet (NCD) (**Figure S2A**). Unexpectedly, under a HFD (60% fat) challenge, male KI^cas9^ mice gained less body weight compared with WT littermates (**Figure 1B**). In agreement with the lean phenotype, KI^cas9^ mice showed an improvement of blood glucose and lipid levels, and lipid storage in the liver (**Figures S2B-H**). KI^cas9^ mice had reduced total fat mass content and less inguinal white adipose tissue (iWAT) (**Figures 1C**, **Figures S2I**). Despite no obvious alterations in food intake, physical activity, or respiratory exchange ratio (RER) (**Figure 1D**, **Figures S2J-K**), KI^cas9^ mice exhibited increased O_2_ consumption (**Figures 1E-F**) and CO_2_ production (**Figures 1G-H**), indicating a higher basal energy expenditure (**Figure 1I**). We further validated the HFD challenge test in the following generation and gained consistent results (**Figure S2L**). In addition, a similar lean phenotype was also observed in female KI^cas9^ mice (**Figures S2M-Q**).

**Figure 1.**
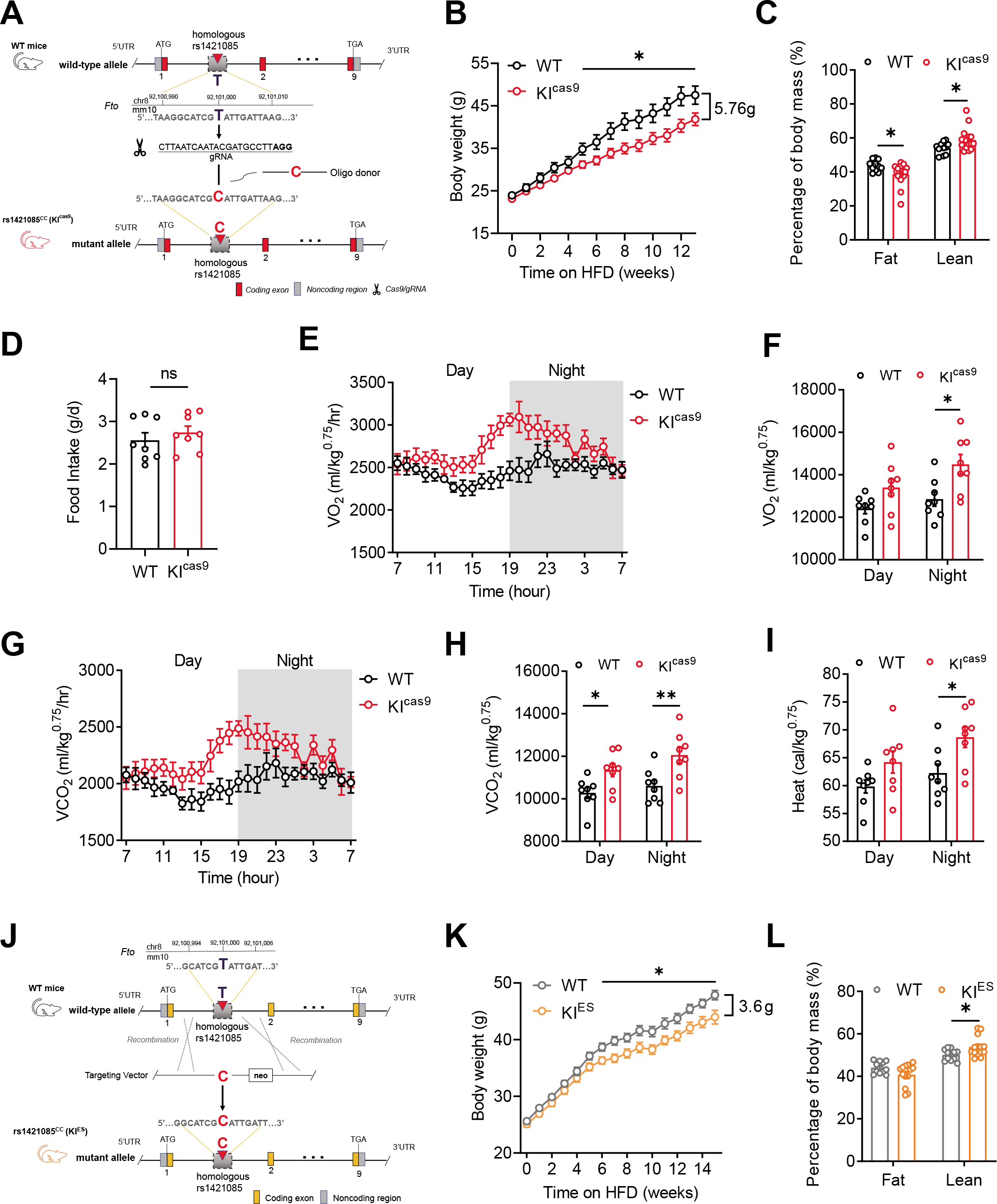
The homologous rs1421085 T>C variant promotes energy expenditure and resists HFD-induced adiposity. **A**, Schematic diagram of the construction of the homologous rs1421085_C knock-in (KI^cas9^) mouse model with CRISPR/Cas9 method. **B-D**, Body weight (**B**), body composition (**C**), and food intake (**D**) of KI^cas9^ mice and WT littermates under HFD challenge initiating from 8-week-old to 21-week-old (B, C, n = 11:16, D, n=8:8). **E-I**, Whole-day and 12h O_2_ consumption rates (**E-F**), CO_2_ production rates (**G-H**), and heat generation (**I**) of KI^cas9^ and WT mice were measured at the end of HFD feeding, with normalization to the body weight (kg^0.75^) (n = 8:8). **J**, Schematic diagram of the construction of the homologous rs1421085_C knock-in (KI^ES^) mouse model with homologous recombination of locus targeting in murine embryonic stem (ES) cells. **K-L**, Body weight (**K**) and body composition (**l**) of KI^ES^ mice and WT littermates under HFD challenge initiating from 8-week-old to 23-week-old (n = 12:13). Data are mean ± s.e.m. of biologically independent samples; unpaired two-sided Student’s *t*-test. * *P* < 0.05, ** *P* < 0.01. ns, not significant.

To further confirm the phenotypes and exclude the possibility of off-targeted editing by CRISPR/Cas9 method, we next utilized the conventional gene targeting strategy with homologous recombination applied to mouse embryonic stem cell (ES) to construct the global rs1421085_CC knock-in (termed KI^ES^) mouse model (**Figure 1J**, **Figure S1C**). Both male and female KI^ES^ mice showed no body weight change under NCD (**Figure S3A-B**) but gained less weight under HFD challenge, with a decreased trend of fat mass and attenuated fat accumulation in liver (**Figures 1K-L**, **Figures S3C-I**). These results suggested that homologous rs1421085 T>C variant resisted HFD-induced adiposity.

### The homologous rs1421085 T>C variant augments the thermogenic capacity of brown adipocytes

Although we observed no significant difference in the morphology of white adipose tissues between KI^cas9^ mice and WT littermates under HFD challenge (**Figure S2H**). Histological analysis indicated that WT mice showed a brown-to-white change in BAT under HFD while KI^cas9^ mice had more condensed and smaller adipocytes in BAT (**Figure 2A**). Consistently, the expression of thermogenesis-related genes, including *Ucp1*, *Pgc-1*α, *Prdm16*, *Cidea*, *Elov6*, *Cox7a* and *Cox8b*, were significantly increased in BAT of KI^cas9^ mice (**Figure 2B**), as were the protein levels of UCP1 and PGC-1α (**Figures 2C-D**). Similar morphological changes and the increased expression patterns of these genes were further validated in BAT of KI^ES^ mice (**Figures 2E-H**, **Figure S3J**), together with a significant increase in mitochondrial quantity (**Figure S3K**). To assess the autonomous effects of homologous rs1421085 T>C variant in brown adipocyte, stromal vascular fraction (SVF) were isolated from BAT of both genotypes and induced to mature brown adipocytes (Wang et al., 2013). An enhanced thermogenic capacity, indicated by increased thermogenesis-related genes (**Figures 2I-K**, **M-O**) and oxygen consumption rate (OCR) (**Figures 2L**, **P**), were observed in mature brown adipocytes derived from both KI^cas9^ and KI^ES^ mice models, when compared with corresponding WT controls. Collectively, these results indicated that homologous rs1421085 T>C variant augmented thermogenesis of brown adipocytes both *in vivo* and *in vitro*.

**Figure 2.**
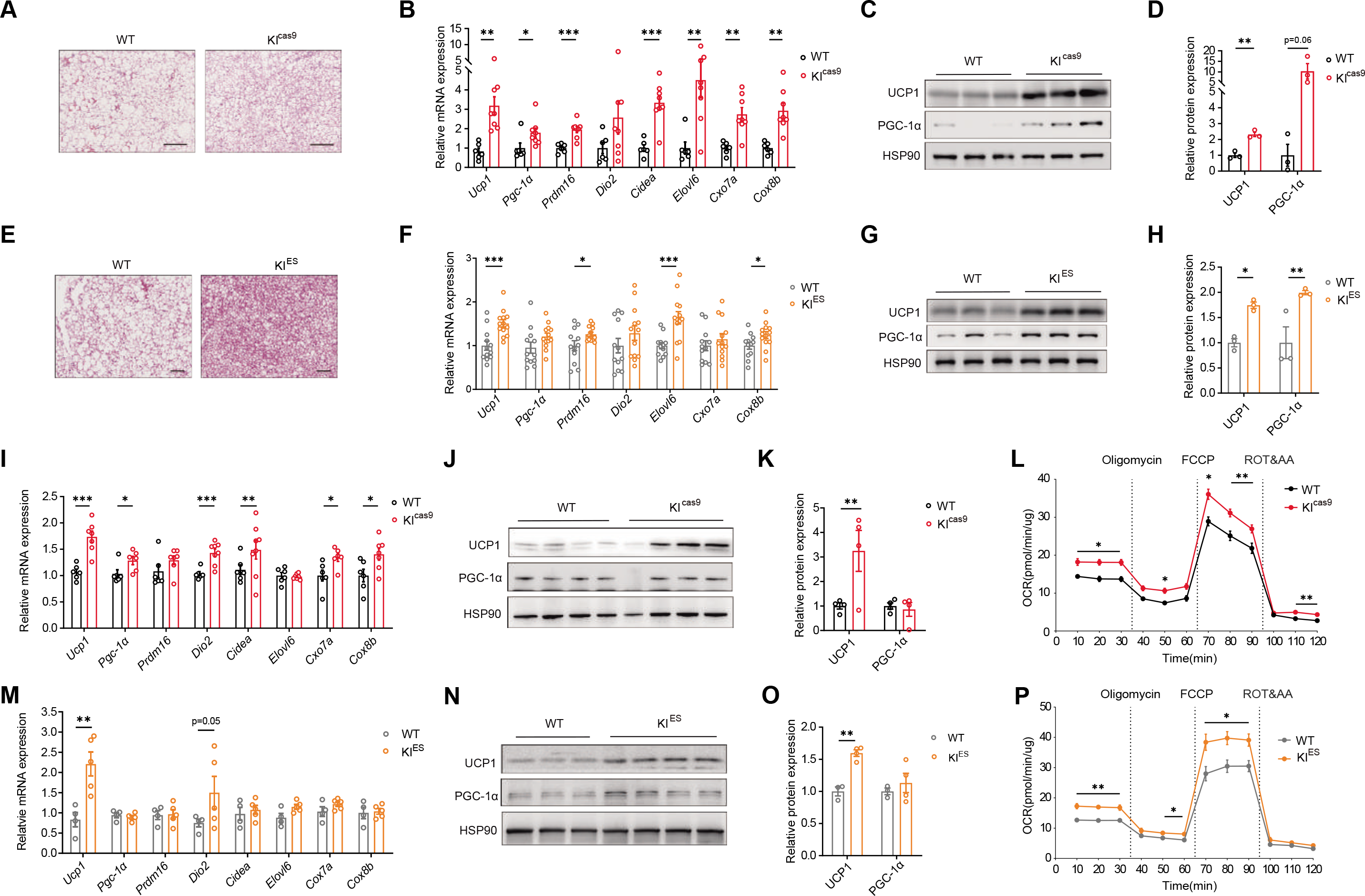
The homologous rs1421085 T>C variant augments thermogenic capacity of brown adipocytes. **A-D**, The representative image of H&E staining (**A**), the mRNA expression of thermogenesis-related genes (**B**), the protein levels (**C**) and the qualification analysis (**D**) of UCP1 and PGC-1α in BAT of KI^cas9^ and WT mice under HFD (**B,** n = 11:16; **C, D**, n = 3:3). **E-H**, The representative image of H&E staining (**E**), the mRNA expression of thermogenesis-related genes (**F**), the protein expression (**G**) and the qualification analysis (**H**) of UCP1 and PGC-1α in BAT of KI^ES^ and WT mice under HFD (**F,** n = 12:13; **G H,** n = 3:3). **I-L**, The mRNA expression of thermogenesis-related genes **(I)**, the protein expression (**J**) and qualification analysis (**K**) of UCP1 and PGC-1α, and oxygen consumption rate (OCR) **(L)** in the induced mature brown adipocytes derived from BAT SVF of KI^cas9^ and WT mice (**I,** n = 4:6; **J, K,** n = 4:4; **L,** n = 4:5). **M-P**, The mRNA expression of thermogenesis-related genes **(M)**, the protein expression (**N**) and qualification analysis (**O**) of UCP1 and PGC-1α, and oxygen consumption rate (OCR) **(P)** in the induced mature brown adipocytes derived from BAT SVF of KI^ES^ and WT mice (**M,** n = 4:6; **N, O,** n = 3:4; **P,** n = 4:5). **A, E**, Scale bar, 100 μm. Data are mean ± s.e.m. of biologically independent samples; unpaired two-sided Student’s *t*-test. * *P* < 0.05, ** *P* < 0.01, *** *P* < 0.001.

### Brown adipocyte-specific knock-in of homologous rs1421085 T>C variant enhances thermogenesis and resists HFD-induced adiposity

To test the hypothesis that homologous rs1421085 T>C variant in BAT might primarily contribute to the obesity-resistant phenotype, we next constructed another brown adipocyte-specific “risk allele” knock-in model by crossing homologous rs1421085^floxp/floxp^ (KI^fl/fl^) mice with *Ucp1*-Cre mice (*Ucp1*-Cre;rs1421085^floxp/floxp^, termed *Ucp1*-KI^fl/fl^) (**Figure 3A**, **Figure S1D**). Firstly, the potential interference effects of the floxp insertion *per se* on the expression of reported targets in KI^fl/fl^ mice were evaluated, and no difference was found as compared to WT littermates (**Figures S4A-E**). Similar to the two global knock-in models, *Ucp1*-KI^fl/fl^ mice showed no obvious change in body weight under NCD (**Figure S4F**), while they displayed a resistance to HFD-induced obesity, indicated by a decrease of body weight and fat mass in comparison with control littermates (**Figures 3B-C**). Smaller lipid droplets and more mitochondria were observed in the BAT of *Ucp1*-KI^fl/fl^ mice, along with a moderate improvement of lipid accumulation in liver (**Figures 3D-E**, **Figure S4G**-**H**).

**Figure 3.**
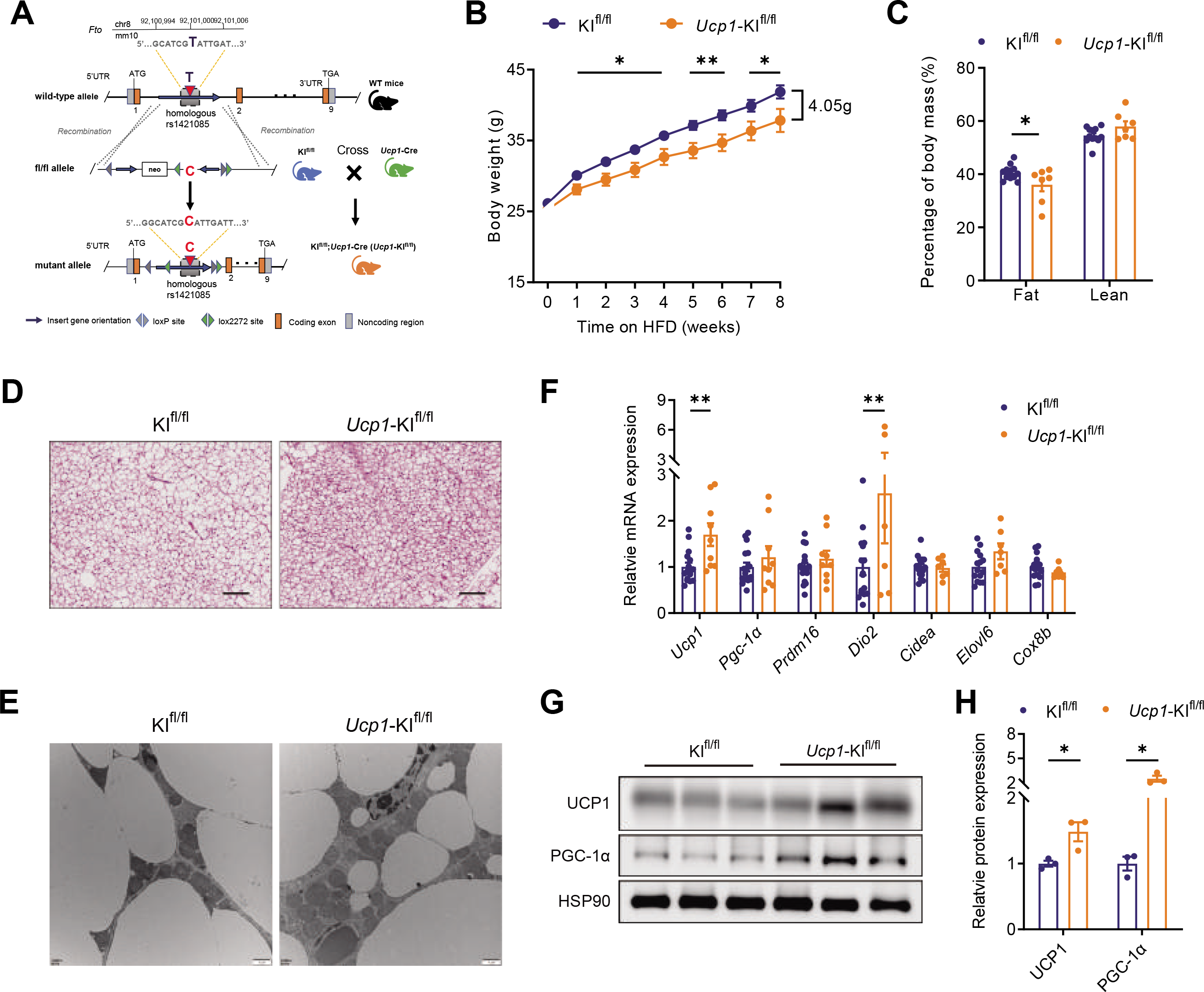
Brown adipocyte-specific knock-in of homologous rs1421085 T>C variant enhances thermogenesis and resists HFD-induced adiposity. **A**, Schematic diagram of the genetic manipulation and construction of homologous rs1421085 variant knock-in mouse model (KI^fl/fl^;*Ucp1*-Cre, termed as *Ucp1*-KI^fl/fl^). **B-C**, Body weight (**B**) and body composition (**C**) in *Ucp1*-KI^fl/fl^ mice and KI^fl/fl^ littermates under HFD challenge initiating from 9-week-old to 17-week-old (**B,** n = 12:7; **C**, n = 8:7). **D-H**, Representative images of H&E staining (**D**) and TEM images (**E**), the mRNA expression of thermogenesis-related genes (**F**), the protein levels (**G**) and quantification analysis (**H**) of UCP1 and PGC-1α in BAT of *Ucp1*-KI mice and KI^fl/fl^ littermates under HFD. **D**, Scale bar, 100 μm; **E,** Scale bar, 1 μm; **D, E, G, H,** n = 3:3; **F,** n = 7:17. Data are mean ± s.e.m. of biologically independent samples; unpaired two-sided Student’s *t*-test. * *P* < 0.05, ** *P* < 0.01.

In consistence, an increased expression of *Ucp1* and other thermogenesis-related genes was further revealed in *Ucp1*-KI^fl/fl^ mice (**Figures 3F-H**). Taken together, with the phenotypes of the two global and one brown adipocyte-specific knock-in models, we illustrated that the biological effect of the homologous rs1421085 T>C variant in BAT contributed prominently to the increased thermogenesis and the resistance to HFD-induced adiposity.

### The homologous rs1421085 T>C variant increases thermogenesis via upregulating *Fto* expression

Several genes neighboring the rs1421085 locus are regarded as effective candidates of the FTO SNP cluster, e.g. *IRX3*/*IRX5*/*IRX6* (Claussnitzer et al., 2015), *RPGRIP1L* (Stratigopoulos et al., 2011, 2016; Tung et al., 2014; Wang et al., 2015), and *FTO* (Grunnet et al., 2009; Klöting et al., 2008), among which *IRX3* and *IRX5* have been shown to increase in subcutaneous preadipocytes of C-allele carriers (Claussnitzer et al., 2015). We unbiasedly and systemically evaluated all the above genes in the BAT of three mouse models, as well as in induced mature brown adipocytes of KI^ES^ and WT genotypes. An obvious increase of *Irx3* expression was detected in KI^cas9^ BAT (**Figure S5A**); increased trends of *Irx3* expression were also observed in other KI models (**Figures S5B-D**); while no significant differences were observed for other candidates (despite increased trends in certain context, like *Rpgrip1l* in KI^ES^ BAT) (**Figures S5A-D**). Unexpectedly, a moderate and sustainable elevation of *Fto* expression was detected in both mRNA and protein levels in each KI model (**Figures 4A-C**). Indeed, the 1 kb genome region containing the “risk allele” of rs1421085 (as well as rs9940128 and rs11642015) showed higher enhancer activity than the major allele (Claussnitzer et al., 2015; Sobreira et al., 2021); however, the downstream target-promoter of these variants has not been clarified. In this regard, four reporter plasmids carrying the 1 kb human genome fragments centered on rs1420185 (T and C allele, respectively) inserted ahead of the human *FTO* promoter (h*FTO*p) sequence either in a forward direction (5’ to 3’) or reverse direction (3’ to 5’), were constructed to investigate the direct enhancer effect of this variant on the activity of *FTO* promoter *per se* (**Figure 4D**). We found that, when linked in a reverse direction, the major allele (T) 1-kb genome fragment enhanced the transcription activity of *FTO* promoter, and the “risk allele (C)” genome fragment further advanced the activity, which did not occur once the fragments were arranged in a forward direction ahead of the promoter (**Figure 4E**).

**Figure 4.**
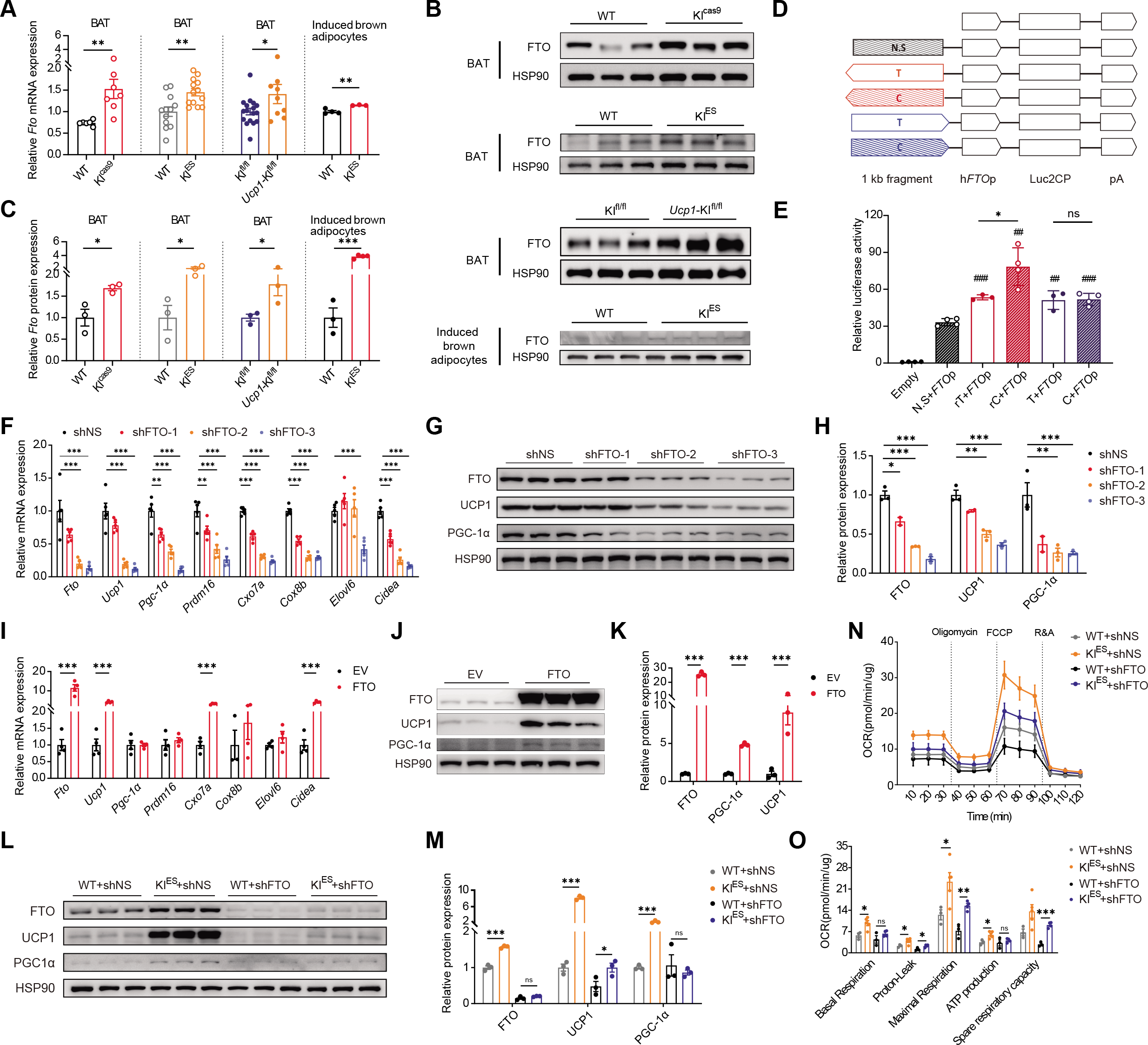
The homologous rs1421085 T>C variant increases thermogenesis via upregulating Fto expression. **A**, The mRNA expression of *Fto* in BAT or mature brown adipocytes of indicated models (KI^cas9^, n = 6:7; KI^ES^, n = 12:13; *Ucp1*-KI^fl/fl^, n = 17:9; induced brown adipocytes derived from KI^ES^, n = 4:3). **B**, **C**, The protein expression (**B**) and quantification analysis (**C**) of FTO in BAT or mature brown adipocytes of indicated models (n = 3:3 in BAT, n=3:4 in induced brown adipocytes). **D, E**, The 1 kb region centered on rs1421085 T>C variation was linked ahead of human *FTO* promoter (h*FTO*p) fragment, and was subsequently fused to *Luc2CP* reporter gene (**D**). Luciferase activity was measured in HEK293T cells transfected with indicated plasmids, in which Renilla luciferase activity (**E**) was used for normalization (n = 3–4). pA, SV40 polyA; ns, not significant. ##, *P<* 0.01; ###, *P* < 0.001 compared with N.S.+ FTOp. **F-H**, The mRNA expression of thermogenesis-related genes (**F**), the protein expression (**G**) and quantification analysis (**H**) of FTO, UCP1 and PGC-1α following *Fto* knockdown in induced mature brown adipocytes derived from BAT SVFs of C57BL/6J mice (MOI = 50; n = 2–4). **I-K**, The mRNA expression of thermogenesis-related genes (**I**), the protein expression (**J**) and quantification analysis of FTO, UCP1 and PGC-1α (**K**) after overexpressing FTO in differentiated SVFs derived from BAT of C57BL/6J mice (MOI = 75; n = 2–4). **L-O**, The protein expression (**L**) and quantification analysis (**M**) of FTO, UCP1 and PGC-1α, as well as OCR (**N**, **O**) after *Fto* knockdown in induced mature brown adipocytes derived from BAT SVFs of wild-type and KI^ES^ mice (MOI = 50; n = 3–4). Data are mean ± s.e.m. of biologically independent samples; unpaired two-sided Student’s *t*-test. * *P* < 0.05, ** *P <* 0.01, *** *P* < 0.001.

The biological roles of FTO in brown adipocytes remains elusive (Wang et al., 2015). We found that, during the differentiation of brown adipocytes *in vitro*, *Fto* expression continuously increased in parallel with the pattern of thermogenesis-related genes such as *Ucp1* and *Pgc-1*α (**Figure S6A**). We also observed a strong positive linear correlation between *Fto* and *Ucp1* mRNA expression (**Figure S6B**). Of importance, *Fto* knock-down remarkably reduced the mRNA and protein expression of *Ucp1* and other thermogenesis-related genes, and *Fto* deficiency impaired the thermogenic capacity of induced brown adipocytes in a dose-dependent manner (**Figures 4F-H**, **Figures S6C-E**); meanwhile, *Fto* overexpression increased their expression (**Figures 4I-K**, **Figure S6F**). Entacapone (ENT), which acts as an m6A demethylation inhibitor of FTO protein, resulted in an obvious impairment of *Ucp1* expression in the induced brown adipocytes (**Figures S6G-J**). These results suggested that FTO protein positively regulated *Ucp1* expression in brown adipocytes.

To better illustrate the roles of FTO in functional BAT pads, and avoid the secondary interference of growth defects observed in global *Fto* knockout mice (McMurray et al., 2013), we crossed *Fto*^floxp/floxp^ mice with *Myf5*-cre mice (Schulz et al., 2013) to effectively delete *Fto* in BAT (*Myf5*-cre;*Fto*^floxp/floxp^, termed MFKO) (**Figure S7A**). Under HFD challenge, MFKO mice showed an increased body weight gain (**Figures S7B**-**C**) and a decline of energy expenditure as compared with control mice **(Figures S7D-K**). Of importance, *ex vivo* experiments showed that UCP1 protein expression decreased upon FTO deletion in the induced brown adipocytes from MFKO mice (**Figures S7L**-**M**).

We further performed RNA immunoprecipitation followed by qPCR (RIP-qPCR) and agarose gel electrophoresis (AGE) tests, and observed that endogenous FTO protein bound to *Ucp1* mRNA in mature brown adipocytes (**Figures S8A-C**). In addition, we also observed an interaction of FTO protein with *C/ebp*α mRNA (**Figures S8B**, **D**), consistent with previous findings in NOMO-1 cells (Su et al., 2020). Moreover, FTO deficiency significantly reduced the stability of *Ucp1* transcripts while did not significantly affect that of *C/ebp*α, suggesting that *Ucp1* was one main target of FTO protein in mature brown adipocytes (**Figures S8E-F**). Importantly, the increased *Ucp1* expression and OCR of KI^ES^ brown adipocytes was largely repressed with either silence of *Fto* gene or enzymic inhibition of FTO protein (**Figures 4L-O**, **Figure S8G-H**). The incomplete elimination of the differences in the tests indicated that other downstream genes of the variant (like *Irx3* (Claussnitzer et al., 2015)) might also be involved in the process (Zhang et al., 2021; Zou et al., 2017). Taken together, these findings revealed that the effects of the homologous rs1421085 T>C variant on the increasing thermogenesis and *Ucp1* expression depended largely on the FTO protein.

### The rs1421085 T>C variant is associated with lower body weight of human infants and its allele frequency parallels with latitudes and ambient temperature of population distribution

Rodents possess thermo-active BAT throughout life, whereas in humans BAT activation peaks after birth to combat the extrauterine coldness for survival, then vanishes with growth (Symonds et al., 2003). To evaluate the potential contribution of BAT degeneration on the adiposity-related outcomes of rs1421085 T>C variant, we surgically removed intrascapular BAT (iBAT) by iBAT excision (ΔiBAT) (Grunewald et al., 2018) in KI^cas9^ and WT littermates (**Figure S9A**). Without iBAT, KI^cas9^ mice lost adiposity resistance under HFD, but appeared to have more body weight increase, higher fat mass percentage, larger white adipocytes and more severe ectopic lipid deposition in the liver when compared to controls (**Figures S9B-F**), in agreement with the adiposity traits of the adult humans carrying the “risk-allele”.

Given the substantial biological effects of homologous rs1421085 T>C variant on murine BAT function and adiposity, we hypothesized that rs1421085 T>C variant may associate with a lower body weight in humans carrying functional BAT. Indeed, a few well-designed longitudinal studies have indicated that the association between *FTO* SNPs and body weight changes dynamically with growth (Sovio et al., 2011). Thus, we performed a meta-analysis to explore the effect of *FTO* polymorphism (including rs1421085 and other three linkage disequilibrium SNPs) on body weight/BMI. In agreement with previous findings (Fang et al., 2010; Hakanen et al., 2009; Lauria et al., 2012; Liu et al., 2010; Mook-Kanamori et al., 2011; Pausova et al., 2009; Scott et al., 2010; Seal et al., 2011; Xi et al., 2010), BMI difference between the effect (minor)-allele and the reference (major)-allele groups emerged since 8 years old; that is, a higher average BMI was observed in the effect-allele group (**Figure S10A**); however, this difference disappeared when the included population was restricted to younger than 8 years old (**Figure S10B**). Importantly, we observed a mild but significant decrease of birth weight in the effect-allele group (P = 0.009, 95% CI: [-0.11, -0.02]; **Figure 5A**). As only a few of studies described the histological and molecular characteristics of human fetal BAT (Velickovic et al., 2014), we collected fetal BAT and confirmed the existence of functional BAT even at middle-gestational stage (**Figure S11**). In further, we found the co-localization of FTO and UCP1 protein in fetal brown adipocytes, which were featured with strong UCP1 and PLIN1 staining (**Figure S11**), suggesting that a potential role of FTO in fetal BAT development and human thermogenesis.

**Figure 5.**
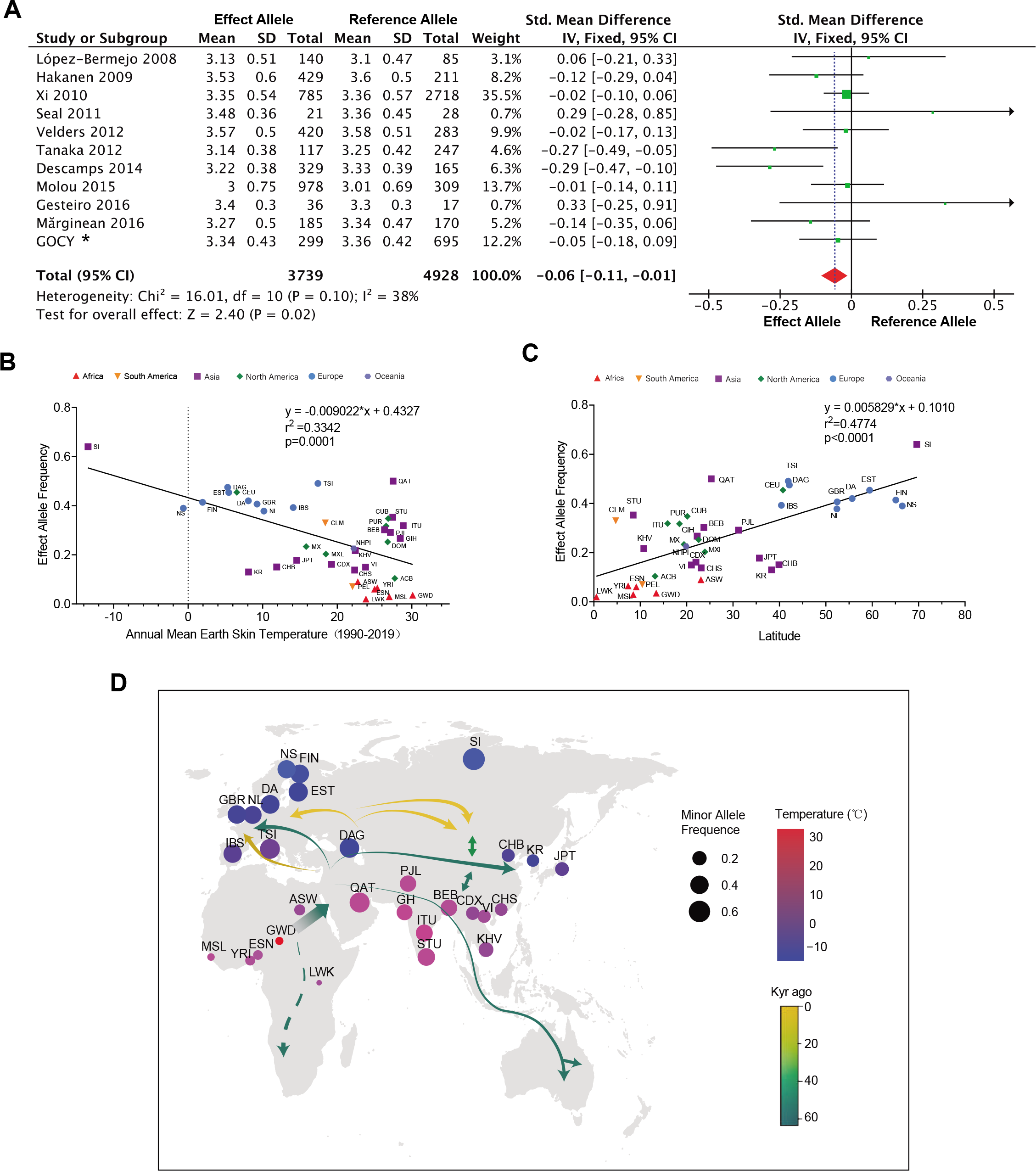
The rs1421085 T>C variant is associated with lower body weight of human infants and its allele frequency parallels with latitudes and ambient temperature of population distribution. **A**, The Forest plot of the association between birth weight and human FTO rs1421085 polymorphism. Studies or subgroups included from previous studies and our in-home data were represented (see Methods). The size of the green box corresponding to each study was proportional to the sample size, while the horizontal line showed the corresponding 95% confidential intervals (CI) of the standard deviation (std) mean difference. The overall std mean differences were based on a fixed-effects model shown by the red diamonds. The solid, vertical line represented the null result. *, data from the Genetics of Obesity in Chinese Youngs (GOCY) study. Effect allele, rs1421085_C; reference allele, rs1421085_T. SD, standard deviation, IV, inverse variance methods. **B**, **C**, Correlation analysis of the effect allele frequency and absolute latitude (**B**) or annual mean earth skin temperature (**C**). Colors and symbols represented populations from different continents. **D**, The relative proportion of the Africa-Eurasia populations carrying rs1421085_C allele. Each circle represented a distinct population: color means earth skin temperature, and size means rs1421085_C allele frequency. Arrows indicated the direction of population migration, and color represents the time of occurrence. **B**-**D**, Population details see **Table S5**.

Although a strong positive association between *FTO* SNPs (including rs1421085) and obesity is confirmed across adult populations of diverse ancestry, a relatively weaker association and a lower frequency of “risk alleles” have been observed according to large-scale GWAS studies based on African populations (Loos and Yeo, 2014; Monda et al., 2013; Peters et al., 2013). To date, reasons for the discrepancy among African and non-African populations remains unclarified. Herein, we assumed that environmental coldness, which activates BAT but rarely occurred in Africa, might contribute to the low frequency distribution of the *FTO* SNPs. To this end, we first analyzed ambient environmental temperatures with the allele frequency of the rs1421085 T>C variant in various populations (of all the 1000 Genomes populations, see **Methods**), and observed a significantly negative correlation in populations settled in Africa-Eurasian continents (*P* = 0.0001, R^2^ = 0.33; **Figure 5B**). The negative correlation remained significant after adjusting Paleozoic temperature (which possibly affected BAT function in ancient humans) using two published models (Eisenberg et al., 2010) (*P* < 0.0001, R^2^ = 0.40, and *P* < 0.0001, R^2^ = 0.39, respectively; **Figures S12A**-**B**). In addition, we analyzed the correlation between the variant frequency and the coordinates latitudes (one determinant factor of ambient environment temperature) of all populations and observed a more robust positive association (*P* < 0.0001, R^2^ = 0.48; **Figure 5C**). Meanwhile, no statistically significant correlation was found between the variant frequency and longitudes or altitudes among populations (**Figures S12C-D**).

Adaption to coldness outside Africa, especially during the last ice age, was a major challenge for early Homo sapiens migrating from East Africa (Clemente et al., 2014).

Acclimatization through genetic accommodation during human migration is a prevailing theory, e.g. the EDAR^V370A^ mutation selected by warm and humid Asia environment (Kamberov et al., 2013), the unique EPAS1 haplotype structure selected by the hypoxic environment of the high-altitude Tibetan platens (Huerta-Sánchez et al., 2014). From this aspect, with modern humans leaving Africa, migrating, and finally settling on other continents, their ambient temperature/residential location shifted from a warm/low latitude to cold/high latitude. While referring to human migration route maps reported by previous studies (Nielsen et al., 2017), we proposed a genetic diffusion map of the rs1421085 T>C variant frequency on the African and the Eurasian continents, that is, the frequency was lowest in East Africa, intermediate in East and South Asia, increased in Europe, and finally reached its zenith in Siberia (**Figure 5D**).

## Discussion

Immediately after the identification of the association between FTO variants and obesity, Palmer’s group made an elegant and well-controlled comparison of the energy balance of FTO “risk allele” carriers versus non-carriers in 4-10 years old children (Cecil et al., 2008). An indirect calorimetry approach was applied to assess basal metabolic rate (BMR), while the doubly labeled water method (used as golden standard (Nagy, 1983) was employed to measure total energy expenditure. Paradoxically, they revealed that the “risk allele” carriers had a higher BMR and total energy expenditure than non-carriers. Nevertheless, despite that covariate adjustment partially abolished the “risk allele” effects on BMR, the total energy expenditure remained significantly different between the two groups (an increase of 1160 kJ in “risk-allele” carriers). Unfortunately, these important findings were ignored as subsequent studies overwhelmingly argued that the “risk allele” had no or a reduction effect on energy expenditure in adults (Claussnitzer et al., 2015; Speakman et al., 2008). The causal effects of the FTO risk allele at the organism level remain to be illusive. It is emphasized that humanized murine models are powerful tools to dissect mechanisms underlying the association of FTO variants with human phenotypes (Herman and Rosen, 2015; O’Rahilly et al., 2016). To our knowledge, to date, there is no report regarding murine models to exactly recapitulate human FTO polymorphisms with single-nucleotide substitution methods. Towards this end, we initially established two strains of global homologous rs1421085 T>C variant knock-in mice using CRISPR/Cas9 and conventional gene-targeting with homologous recombination strategies, respectively. A third strain of brown adipocyte-specific homologous C-allele single-point knock-in mice was constructed to confirm its biological roles in BAT. In support of Palmer’s findings in children (Cecil et al., 2008), our study reported the FTO variant yielded an enhancement of thermogenic gene expression, oxygen consumption, and total energy expenditure of brown adipocytes both *in vivo* and *ex vivo*. Another important finding is that the rs1421085 T>C variant is indeed a pivotal genetic trigger modulating a line of downstream genes, among which *Fto* showed the most stably increased expression in “risk-allele” group. Furthermore, we provided a reliable evidence of FTO protein directly regulating *Ucp1* expression both *in vivo* and *in vitro*, which was consistent with the obese phenotype observed in adipocyte-specific *Fto* knockout mice (Wang et al., 2015), pointing out the involvement of FTO in the function of the rs1421085 T>C variant. Except for *Fto,* we also validated an increased expression of *Irx3*, *Irx5* (Claussnitzer et al., 2015; Smemo et al., 2014; Sobreira et al., 2021) and *Rpgrip1l* (Stratigopoulos et al., 2016) in certain contexts, supporting a temporal, spatial, and cell type-dependent regulation by the variant (Sobreira et al., 2021). Furthermore, we have previously revealed that IRX3 overexpression promoted the transcriptional activity of *Ucp1* and enhanced thermogenesis in brown adipocytes (Zhang et al., 2021; Zou et al., 2017), which might be one plausible explanation for the incomplete rescue of FTO knockdown or inhibition to the rs1421085 T>C variant’s effects. We want to mention that these findings in the knock-in models of this single-nucleotide variant is not completely the same as in the recently reported two mouse models harboring a deletion of 82 bp and 20 kb containing the same FTO variant, respectively (Laber et al., 2021; Sobreira et al., 2021), which may be due to non-specific deletion of other potential functional motifs in the region by their strategies. Our models that recapitulate the same human variant are more specific for the evaluation of the pathophysiological changes produced by the homologous rs1421085 T>C variation at the organism level.

The first available document about BAT deposition in infants was reported by Neumann in 1902 (Ravussin and E.Galgani, 2011). BAT in human newborn infants produces heat by non-shivering thermogenesis to maintain body temperature. When the infants were shifted from 34–35c:: (thermoneutral temperature) to 25–26c:: (cooling temperature), their rectal temperature dropped 0.1–0.5c:: and O_2_ consumption almost doubled, accompanied by a relatively warmer skin temperature of the intrascapular BAT region than other sites (Dawkins and Scopes, 1965). BAT depletion or BAT functional defects are associated with the development of “cold syndrome”, which threatens newborn infants’ survival (Marsh, 1983). These phenomena gave rise to speculation that higher BAT content or activity would increase the survival of infants born in ambient cold temperatures. For example, the Last Glacial Maximum (LGM), when the mean temperature was 6.1c:: lower than nowadays (Tierney et al., 2020), challenged all species including modern humans. Some species adapted to the prehistoric climate successfully (Kumar et al., 2015), while others, e.g., archaic humans, *Neanderthals*, *Denisovans*, and *Homo erectus*, became extinct (Stewart and Stringer, 2012). To date, factors that determine such successful adaptation are still poorly understood (Dibble et al., 2017; Enerbäck, 2010). We found that rs1421085 T>C knock-in increased the thermogenic capacity of mice. In this context, the functional FTO variant–rs1421085 T>C–has likely increased the survival odds of human newborn infants by increasing their thermogenesis. In consistent with this hypothesis, we found that the frequency of rs1421085 T>C was the lowest in East and Middle Africa, higher in East Europe, much higher in North Europe, and reached the highest in Siberia. Interestingly, the association between the FTO variants and obesity in European and Asian populations is not well-validated in Africans, and only mild effect was observed in a few studies (Loos and Yeo, 2014; Monda et al., 2013; Peters et al., 2013). Furthermore, a tight linkage disequilibrium among various obesity-related GWAS positive signals that clustered within the first intron of the *FTO* gene is observed in non-African populations, while this genetic architecture of the FTO loci breaks down in native Africans (Peters et al., 2013). These findings suggest that this genomic region has experienced genomic recombination and undergone genetic selection since modern humans descended from several branches of Homo sapiens populations that migrated out of Africa (Loos and Yeo, 2014). Several possibilities exist regarding the origin and distinct distribution frequency of the rs1421085 T>C variant between Africans and non-Africans. One hypothesis is that the variant arose occasionally in African populations, and after encountering extremely cold temperatures during the migration away from Africa, those newborns carrying the variant gained higher thermogenic capacity to thrive. The frequency subsequently increased stepwise, in parallel with migration, which started in the hot/low latitudes and gradually increased when moving to the cold/high latitudes. Another hypothesis claims that, in cold temperatures, an intense genomic recombination within the first intron of the *FTO* gene induced random occurrence of rs1421085 T>C in a variety of populations, independently. Populations in Africa rarely encountered such cold exposure, thus the variant was merely observed. According to this hypothesis, the rs1421085 T>C variant first presented in archaic humans after they settled in Eurasia, later the genomic fragment containing the variant possibly flowed back to Africa. This process might have happened repeatedly (Henn et al., 2012). It should be pointed out that all these possibilities warrant further genetic evidence of modern and archaic humans for confirmation or exclusion. Of note, a few nucleotide sequences centered to the rs1421085 locus were reported to be transcription factor (e.g., ARID5B and CUX1 (Claussnitzer et al., 2015; Stratigopoulos et al., 2016)-binding elements, showing a high conservation among species, even including zebrafish (Claussnitzer et al., 2015). These results indicate the *cis*-elements might be essential for the expression of downstream targets, while our data support that the T>C alteration belongs to a gain-of-function mutation. In future studies, element disruption at the locus would be informative for us to understand the roles of wild-type genotypes in obesity and other pleiotropic outcomes. Despite that BAT brought evolutionary advantages for mammals to survive cold stress (Cannon and Nedergaard, 2004; Smith and Roberts, 1964), whether the thermogenic variation contributes to natural selection in the evolution of mammalian endothermy is also a matter of uncertainty (Jastroch et al., 2018). It should be noted that, in additional to the rs1421085 T>C variant, other FTO variants also have functional impacts on the target genes (Sobreira et al., 2021; Stratigopoulos et al., 2016). The various haplotypes at convergence of different variants would lead to fluctuation of the phenotypes, including energy expenditure and appetite (Harbron et al., 2014).

On the other hand, how to re-evaluate the predisposition of the “risk-allele” to adiposity in adult humans? According to the Early Growth Genetics (EGG) Consortium data, a shift pattern of the effect of the *FTO* rs9939609 T>A variant on BMI was observed in adolescents of different ages: with a positive association between the “risk allele” and BMI from 5.5 years old onwards; however, there was an inverse association below 2.5 years old (Sovio et al., 2011). In consistence, the genetic influence of the same variant (rs9939609) on BMI was progressively stronger with increasing age (from 0.48 at 4y to 0.78 at 11y), as reported from a longitudinal large twin cohort of more than 7,000 children (Haworth et al., 2008), which further indicated an age-related characteristic of FTO variants in the regulation of adiposity. In humans, BAT diminishes rapidly in the first decade of life (Heaton, 1972; Lean et al., 1986). To test whether this disappearance influence the biological effect of *Fto* variant on adiposity phenotypes, we removed intrascapular BAT of both KI^ES^ and WT mice, and successfully simulated the predisposition of the “risk allele” to obesity, as observed in humans. This interesting finding provided some potential underlying explanations for the phenotype shift from obesity-resistance to obesity-sensitivity of the rs1421085 T>C variant, such as the compensatory role of this variant in other tissues in addition to BAT (e.g., hypothalamus and WAT), the reduction of certain metabolic-protective secretory factors (e.g., BMP8b, FGF21 and IL-6) of BAT (Villarroya et al., 2017), or the long-term consequence of altered feeding/eating habits at early life and following catch-up growth (Reilly et al., 2005). In a further speculation, other potential downstream targets of rs1421084 T>C, e.g., *IRX3*/*IRX5*/*IRX6* (Claussnitzer et al., 2015; Sobreira et al., 2021) and *RPGRIP1L* (Stratigopoulos et al., 2016), as well as the target of *FTO per se* (Li et al., 2017), could also participate in the compensatory predisposition to adiposity.

## Conclusion

In summary, by collecting evidence from population studies as well as genetically modified murine models, we uncovered the possible biological roles and molecular basis of the rs1421085 T>C variant and postulated the rs1421085 T>C–*Fto*–*Ucp1* axis in the regulation of BAT thermogenesis. These findings may enhance our understanding regarding the possible “causative biological functions” of the “risk variants” derived from extensive GWAS data especially for those localized in non-coding regions and further remind us of the importance of genome-environment interaction, and the potential contribution of genetic variants in the process of adaptive evolution.

## Acknowledgments

We thank all the participants for their involvements in this study, and we thank Dr. Juan Shen and Dr. Qijun Liao from BGI-Shenzhen for their suggestive comments on adaptive evolution of the FTO variant.

## Author contributions

J.W. designed the experiments and supervised the study. Z.Z., N.C., N.Y., and Y.H. carried out the animal and molecular experiments. J.W., N.Y., and M.T. analyzed the genetic data. J.W. and R.L. analyzed all the experimental data. D.L, A.G., P.L., and H.L. provided supports in animal model tests. G.N., W.W., J.H., W.G.., and W.W. provided the cohort resources. J.W., R.L., and Z.Z. wrote the manuscript. G.N. and L.Q. contributed to text revision and discussion.

## Methods

### Mice

The first homologous rs1421085 T>C variant knock-in mouse line was generated through CRISPR/Cas9 technology (Jinek et al., 2012), termed as KI^cas9^. In brief, a single-guide RNA (sgRNA) sequence (sgRNA 5′-CTTAATCAATACGATGCCTTagg) was selected to overlap amino acid residue T41 in murine *Fto*. Cas9 mRNA, sgRNA, and donor oligo were co-injected into zygotes to produce the homologous rs1421085_C genotype mouse. The GGCATCG***T***ATTGATT to GGCATCG***C***ATTGATT mutation site in donor oligo was introduced into intron 1 of murine *Fto* locus by homology-directed repair and verified by PCR and Sanger sequencing. Among the predicted potential off-target sites top 10 reported previously with CCTop software (Stemmer et al., 2015), none were reported for an obesity-related phenotype (**Table S1**). In addition, Sanger sequencing of a 6 kb region centered on the homologous rs1421085 locus revealed no other mutations (**Figure S13**). F1 heterozygous targeted mice were self-crossed to generate the F2 homozygous KI^cas9^ mice, and WT littermates were used as controls. The second homologous rs1421085 T>C variant knock-in mouse line was generated by the general targeting strategy of homologues recombinant in embryonic stem cell (ES), termed as KI^ES^. In brief, the c.45+60752 T>C point mutation was introduced into mouse *Fto* intron 1 in the 5’ homology arm to construct a linearized vector, which was then transfected into C57BL/6 ES cells according to standard electroporation procedures. The transfected ES cells were selected and picked for DNA isolation. Subsequent PCR screening and Sanger sequencing were performed to select positive homologous recombination, and correctly targeted clones were further confirmed by Southern blot analysis. Targeted ES cells were micro-injected into C57BL/6 albino embryos and then re-implanted into CD-1 pseudo-pregnant females. Germline transmission was confirmed by breeding with C57BL/6 females and subsequent genotyping of the offsprings. F1 heterozygous targeted mice were confirmed by PCR and Sanger sequencing, then self-crossed to generate F2 homozygous KI^ES^ mice, and WT littermates were used as controls. To generate conditional knock-in mice carrying the homologous rs1421085 T>C variant, the mutant partial intron 1 of the murine *Fto* gene was cloned upstream of exon 2 in the reverse orientation. The mutant partial intron 1 and corresponding WT fragment were flanked with lox2272 and loxP (**Figure 3A**), respectively. The F1 heterozygous targeted mice were confirmed by PCR and self-crossed to generate F2 homozygous mice carrying the floxed homologous rs1421085 T>C variant that were termed KI^fl/fl^. *Ucp1*-Cre mice were obtained from the Jackson Laboratory (Jax no. 024670) and crossed with KI^fl/fl^ mice to breed KI^fl/fl^;*Ucp1*-Cre offsprings (termed as *Ucp1*-KI^fl/fl^), with KI^fl/fl^ littermates as controls. *Fto*^fl/fl^ mice (Jax no. 027830) and *Myf5*-Cre (Jax no. 007893) were cross-linked to generate *Fto*^fl/fl^;*Myf5*-Cre (termed as MFKO) mice, with *Fto*^fl/fl^ littermates as controls.

Genomic DNA from the tail of the newborn mice was extracted within 3-weeks of birth. The KI^ES^ and KI^cas9^ knock-in mice were verified by DNA sequencing, while the *Ucp1*-KI^fl/fl^ knock-in mice were verified by PCR and DNA sequencing. All primers used for genotyping were listed in **Table S2**. For stromal vascular fraction (SVF) isolation, wild-type C57BL/6J mice aged 4–6 weeks were obtained from Lingchang Biotech (Shanghai, China). Mice were housed in standard cages in a specific-pathogen-free facility on a 12-h light/dark cycle with *ad libitum* access to food and water. For HFD test, same-sex littermates were housed in groups of up to four mice per cage and fed a 60 kcal% HFD diet (Research Diet 12492i). For NCD feeding, same-sex littermates were housed in groups of up to five mice per cage. Average food intake was measured in each cage for three to four days and was replicated thrice. All animal procedures were approved by the Animal Care Committee of Shanghai Jiao Tong University School of Medicine (SJTUSM) and followed the guide for the care and use of laboratory animals.

### IPGTT, ITT and lipid measurement

For the intraperitoneal glucose-tolerance tests (IPGTT), mice were fasted overnight and injected with glucose (1.5 g/kg body weight) intraperitoneally (i.p.). For the insulin tolerance test (ITT), mice were fasted for 6 h and injected i.p. with recombinant human insulin (1.5 U/kg, Eli Lilly). Blood glucose was measured at 0, 15, 30, 60, 90 and 120 min, respectively, after administration using a Contour next glucometer (Bayer). Plasma triglyceride (TG), total cholesterol (TC), low-density lipoprotein-cholesterol (LDL-c) and high-density lipoprotein-cholesterol (HDL-c) were measured with commercial kits according to the manufacture’s guidance (Kehua Bioengineering, Shanghai, China).

### Body composition analysis and measurement of whole-body energy expenditure

Body composition was measured using an EchoMRI-100H (EchoMRI, USA). For the evaluation of whole-body energy expenditure, the mice were placed in a Comprehensive Laboratory Animal Monitoring System (CLAMS, Columbus Instruments, USA), individually in a single cage. O_2_ consumption, CO_2_ output and physical activity were continuously measured three to four times per hour, lasting for 48 h. The respiratory exchange ratio (RER) and heat generation were calculated based on the O_2_ and CO_2_ data. All data of O_2_ consumption, CO_2_ production and heat generation were normalized to (body weight)^0.75^ (Tschöp et al., 2011).

### Morphological analysis

For hematoxylin and eosin (H&E) staining, the adipose and liver tissues were isolated, fixed in 4% paraformaldehyde, embedded in paraffin, and sliced into 5-μm for WAT and liver, and 3-μm for BAT sections, respectively. High resolution images were captured under a microscope (Olympus, Japan), while global pictures were scanned using a Digital Pathology Slide Scanner (KFPRO-120). For transmission electron microscopy (TEM), fresh BAT samples were immediately pre-fixed in 2.5% glutaraldehyde PB for 2 h, rinsed twice with PB buffer for 10 min each time, fixed with 1% osmium acid PB for 2 h, and rinsed a further twice with double distilled water for 10 min each. After stepwise dehydration and propylene oxide replacement, tissues were then embedded using Epon812 solution and propylene oxide, and subsequently heated in a 60 c:: oven for 48 h. Ultra-thin sections were obtained using an EM UC7 (Leica, Germany) and these were stained with lead citrate. Electron micrographs were imaged under H-7650 transmission electron microscope (Hitachi, Japan).

### Immunofluorescence analysis

For co-immunoprecipitation (IF), the paraffin section of human embryonic brown adipose samples was successively stained with UCP1 antibody (Abcam, ab10983; 1:500), Perilipin antibody (CST, 9349s; 1:200) and FTO antibody (Abcam, ab126605; 1:300) using Tyramide Signal Amplification (TSA) strategy as described (Hermsen et al., 2021). The try-488, try-cy3 and try-cy5 were used as tyramine conversion reagents. The 1mmol Tris-EDTA (PH=9.0, Sigma) was used for Thermoantigen repair. Antifade Mounting Medium with DAPI (Beyotime) was used for sealing. Automated acquisition system (TissueFAXS Plus, TissueGnostics GmbH, Austria) was used for whole slide scan.

### RNA extraction and real-time PCR analysis

Total RNA was extracted from cultured cells or frozen adipose tissue using the Eastep Super Total RNA Extraction Kit (Promega (Beijing) Biotech Co., China). The absorbance ratio at 260/280 nm and the RNA concentration of each sample were detected using a NanoDrop ND2000 spectrophotometer (Thermo Scientific). Reverse transcription was performed using the PrimeScript Reverse Transcript Master Mix (TaKaRa, Japan). Quantitative PCR was performed using a QuantStudio Dx Real-Time PCR Instrument (Applied Biosystems Instrument). The comparative 2^-ΔΔCT^ method was used to evaluate the relative mRNA levels; *36B4* served as the reference gene. Primers used for PCR are listed in **Table S2**.

### Protein preparation and Western blotting analysis

Total protein was isolated using RIPA lysis buffer (Biocolors, China) with a protease inhibitor cocktail (Sigma, USA). All protein samples were subjected to concentration determination and immunoblotting assay with the indicated antibodies. HSP90 (4877s, CST, USA) was used as the internal control, and the following primary antibodies were used: anti-FTO (ab92821, Abcam, USA; 45980S, CST, USA), anti-UCP1 (ab10983, Abcam, USA), and anti-PGC-1α (sc514380, Santa, UAS).

### Plasmid construction and luciferase reporter assay

Human *FTO* promoter (h*FTO*p, -1000 ∼ +51 bp from transcriptional start site) was amplified from human genomic DNA (NCBI Reference Sequence: NC_000016.10). For the blank control, h*FTO*p was inserted into the pGL4.12 vector (Promega) between the XhoI and HindIII restriction sites. For enhancer activity analysis, the 1 kb fragment including human FTO SNP rs1421085_T (the reference allele) and rs1421085_C (the minor allele) were constructed using the QuikChange II site-directed mutagenesis kit (Stratagene) and inserted in forward or reverse orientation ahead of h*FTO*p into the pGL4.12 vector, respectively (**Figure 4D**). All constructs were verified by DNA sequencing. HEK293T cells were purchased from ATCC (CRL-11268). Cells were seeded in 24-well plates in DMEM supplemented with 10% fetal bovine serum (FBS, Gibco) before transfection. After a 24 h transfection with 1000 ng h*FTO*p reporter plasmid along with 1 ng pRL-SV40 plasmid, HEK293T cells were harvested for luciferase activity assessment using a dual-luciferase reporter assay system (Promega). The results were normalized to Renilla luciferase activity.

### Lentiviruses production and infection

Lentiviral mouse *Fto* shRNA (shFTO1-3) constructs targeting mouse NM_011936 were assembled using pLKD-CMV-G&PR-U6-shRNA lentivector cloning. The siRNA sequences used were listed in **Table S3**. The shRNA sequences were inserted between the *AgeI* and *EcoRI* restriction sites. Human FTO overexpression plasmid was constructed with the pLenti-EF1a-EGFP-P2A-Puro-CMV-MCS-3Flag vector. The 1509 bp fragment of human FTO coding sequence was inserted between EcoRI restriction sites. All constructs were verified by DNA sequencing. The shRNA and primers are listed in **Table S3**. The SVFs were infected with the lentivirus 12 h after seeding at a multiplicity of infection (MOI) ranging from 50 to 100 according to the manufacturer’s instructions. Polybrene (5 μg/ml) was used to optimized infection efficiency. The SVF medium was changed to culture medium or induction cocktail within 48 h after infection.

### Brown adipocyte differentiation *in vitro*

The SVFs were isolated from the BAT of male mice aged 3-4 weeks and then induced to fully differentiated mature brown adipocytes as previously described(Zou et al., 2017). In brief, the fat pads were isolated, cut into pieces, and fully digested with type II collagenase (Sigma) at 37 °C for 40-50 min, followed by quenching with DMEM/F12 supplemented with 10% FBS. The suspended samples were filtered using a 40-mm strainer (BD, USA) and then plated on 6 cm culture dishes. The SVFs were first grown to 100% confluence in DMEM/F12 supplemented with 10% FBS (plus 1% penicillin/streptomycin and 1 mM L-glutamine), then differentiated into mature brown adipocytes in a cocktail containing 5 mg/ml insulin (Eli Lilly, USA), 1 mM dexamethasone, 1 mM rosiglitazone, 1 mM triiodothyronine (T3), and 0.5 mM IBMX (all Sigma–Aldrich, USA) for the first two days, and subsequently in medium with insulin, rosiglitazone, and T3 for day 3 to 6, depending on experimental requests (see results for details).

### Oil Red O staining

After eight days of induction, mature adipocytes were prepared for Oil Red O staining. In brief, the cells were fixed in 4% paraformaldehyde for 30 min, rinsed with PBS several times, then left to stand at room temperature for 2–4 hours until air-dried. They were subsequently incubated with Oil Red O (Nanjing Jiancheng Bioengineering Institute, China) for 30 min. Images were captured under a light microscope (Olympus).

### Brown adipocyte respiration

SVFs were seeded in an XF24 V28 microplate (Agilent Technologies, USA) coated with poly-L-lysine, and induced to mature brown adipocytes as previously mentioned. The oxygen consumption rate (OCR) was measured on the fourth day of induction using an XF24 analyzer (Agilent Technologies) following the manufacturer’s instructions. Briefly, the induced brown adipocytes were washed with Seahorse assay medium, consisting of XF DMEM supplemented with 10 mM XF glucose, 1 mM XF pyruvate, and 2 mM XF L-glutamine, followed by incubation with 525 ml of assay medium at 37°C in an incubator without CO_2_ (Agilent Technologies) for 1 h. Respiratory inhibitors (75 ml) were loaded into the injection port with the final concentrations of 1 mg/ml oligomycin, 2 mM FCCP, 0.5 mM antimycin A and 0.5 mg/ml rotenone to detect uncoupled respiration, maximal respiration, and nonmitochondrial respiration, respectively. The final OCR results were standardized to the total protein content per well.

### RNA stability assay and methylation quantification

Induced mature brown adipocytes upon *Fto* knockdown were treated with 5 mg/mL actinomycin D (A9415, Sigma-Aldrich) to assess RNA stability of *Ucp1* and *C/ebp*α transcripts. Cells were harvested at indicated time points, and total RNA was extracted by reverse transcription and qPCR analysis as described previously. The half-life (t1/2) of mRNA was determined as the reference(Rui Su, Lei Dong, Chenying Li, …, Jie Jin, Chuan He, 2018). Total m6A levels of the extracted RNA were measured using an EpiQuik m^6^A RNA Methylation Quantification Kit (Colorimetric; Epigentek, New York, USA) according to the manufacturer’s instruction. Samples of 200 ng RNA of each sample was used for assessment.

### RNA immunoprecipitation (RIP) analysis

Induced mature brown adipocyte pellets were washed with PBS and lysed with NET2 buffer (50mMTris, pH 7.4; 150 mM NaCl; 0.1% NP-40, 1:100 PMSF) at 4□::. Aliquotes of cell lysate (40 μl) were saved as input (blank control) for Western blotting and reverse transcription-PCR, respectively. The remaining lysates were equally divided and incubated with 7 μl IgG (negative control, CST) and FTO antibody (Abcam), respectively, for 2 h at 4□::. Magnetic beads (Pierce™ Protein A/G Magnetic Beads) were washed thrice with NET2 buffer, then incubated with cell lysate-antibody conjugation for 2 h at 4□::. The protein-captured beads were washed three times and re-suspended in 1ml NET2 buffer. A total of 200 μl of FTO antibody-conjugated beads was denatured for Western blotting assay to confirm IP efficiency. Remaining beads were eluted in wash buffer containing proteinase K (Thermo) and incubated at 37 □:: for 1 h. RNA from the beads was extracted using Phenol-Chloroform-Isoamylol. The fold enrichment was detected by qPCR as described before, and further confirmed with reverse transcription-PCR and subsequent agarose gel (2%) electrophoresis.

### Fetal brown adipose samples

Human fetal brown adipose samples were obtained from two fetuses aged 20–24 weeks gestation after legal medical termination due to developmental malformation or severe hereditary diseases. Located in interscapular subcutaneous region, two autopsy samples that had tough and firm texture (sand-like granules), and relatively deep color compared with surrounding adipose tissues were included for immunofluorescent staining. The gestational age was estimated by crown-rump length and maternal history. The tissues were collected under informed consent from the parents after the decision to abort was finalized, and the protocol was conducted in compliance with the guidelines and approved by the Medical Ethics Committee of Shengjing Hospital of China Medical University.

### Meta-analysis

We searched for research articles published before October 2020 in PubMed, Web of Science and Embase. To explore the association between FTO polymorphism and BMI in juveniles, the following keywords were used: (Infant or Newborns or Child or Children or Adolescents or Teens or Teenagers or Youths or Childhood) and (rs9939609 or rs8050136 or rs1421085 or rs17817449 or rs1121980 or FTO or FTO protein) and (BMI or Body weight). For the analysis of FTO and birth weight, the following keywords were used: (rs9939609 or rs8050136 or rs1421085 or rs17817449 or rs1121980 or FTO or FTO protein) and (Birth or Birth weight). If the same case series were used in more than one research article, only the one with the largest sample size was included. Inclusion criteria: (1) accessible data of population BMI or birth weight; (2) using case-control, cross-sectional, or cohort design; (3) statistics performed according to genotypes. The following information was extracted: (1) name of the first author; (2) year of publication; (3) country of origin; (4) ethnicity of the study population; (5) sample size; (6) average age, BMI and/or birth weight of each cohort; (7) adjusted covariates; (8) SNP included; (9) Hardy-Weinberg equilibrium (HWE) test values. Exclusion criteria: (1) research with incomplete data; (2) basic research; (3) conference reports and review literature; (4) repeated publications. Details of all studies included are listed in **Table S4**. In addition, the data from the Genetics of Obesity in Chinese Youngs (GOCY) study, which was previously established by Ruijin Hospital and registered in ClinicalTrials.gov (ClinicalTrials reg. no. NCT01084967, http://www.clinicaltrials.gov/) (Chen et al., 2020), were also included to integrate Chinese newborn’s information. Birth weight data were obtained from birth certificates provided by parents, as describe previously (Hong et al., 2013). Genomic DNA was extracted from peripheral blood leukocytes with a QIAmp blood kit (Qiagen, Chatsworth, CA, USA), and rs1421085 variant was testified using a LightCycler480 Taqman PCR system (Roche Diagnostics Ltd, Rotkreuz, Switzerland).

The GOCY study was approved by the Institutional Review Board of the Ruijin Hospital, SJTUSM and was performed in accordance with the principle of the Helsinki Declaration II. A written informed consent was obtained from each participant. RevMan 5.3 software was used for Meta-analysis. For continuous data (such as BMI and birth weight), the standardized mean difference (SMD) was used for effect evaluation, and the 95% confidence interval (CI) was calculated. Various studies were tested for heterogeneity. A fixed-effects model was adopted if P value > 0.05, I^2^ < 50% (small heterogeneity). A random effects model was adopted if P value ≤ 0.05, I ≥ 50% (high heterogeneity). Test level was defined as α = 0.05.

### Variant frequency analysis

Data from a total of 39 distinct ethnic populations living on the African and Eurasian continents were included, among which 26 were extracted from the 1000 Genomes Project with the effect allele frequency obtained from the online database (https://www.internationalgenome.org). The other 13 populations were extracted from different cohorts with gene frequency data derived from NCBI (https://www.ncbi.nlm.nih.gov/snp/rs1421085#frequency_tab). Full names corresponding to the abbreviation of the populations are listed in **Table S5**. The coordinates of populations were obtained based on the characteristics of population migration. For native populations, the coordinates of the cities (with the sampling locations clearly given on the official website of the 1000 Genomes Project) were selected. For the populations with different origins and current residences, the coordinates of the origins were selected. For the populations with unclear origin or that were widely distributed, the coordinates of the country’s capital or the geographical center of the regions were selected. The longitude and latitude of each coordinate were identified using a Google satellite map. The altitude and the annual ambient temperature (30-y mean) of each coordinate was obtained from the official website of NASA (https://power.larc.nasa.gov/data-access-viewer/). Two temperature-corrected models were used to simulate the Paleozoic climate to reduce the bias of the unmatched occurrence of SNP selection (occurred thousands of years ago) and ambient temperature(Eisenberg et al., 2010).

### Statistical analysis

Data were assessed for normal distribution before performing two-tailed, unpaired Student *t*-tests and plotting the figures as mean ± sem. Spearman’s correlation analysis was performed for associations analysis. A P value < 0.05 was considered significantly different. For molecular experiments, data were generated from at least three independent experiments. The SAS statistical system (v.8.0; SAS Institute, Cary, NC, USA) was used for data analysis. A topographic map of populations with different genetic variant frequency was drawn with R (v.3.6.2) and open-source R packages ggplot2, ggmap, RgoogleMaps, sp, maptools, maps, and ggThemeAssist.

Abbreviation and corresponding full-name used in this study are listed in **Table S6.**

## Legends for Supplemental information

**Figure S1 | Sequencing of the homologous rs1421085 T>C variant in human and mouse.**

**A**, Conserved motif module surrounding rs1421085 locus in human and mouse.

**B-C**, Sanger sequence analysis of KI^cas9^ (**B**) and KI^ES^ (**C**) mice, respectively.

**D,** Floxp (left, top) and Cre expression (left, bottom) and BAT DNA sequence analysis (right) of homologous rs1421085 mutation in KI^fl/fl^ and Ucp1-KI^fl/fl^ mice.

**A-D**, Gene sequencing alignment was analyzed with Jalview Version 2 (see **Methods**).

**Figure S2 | The homologous rs1421085 T>C variant protests HFD-induced obesity but shows no change in body weight under chow diet in KI^cas9^ mice.**

**A**, Body weight curve of KI^cas9^ mice and WT littermates under normal chow diet (NCD) (n = 10:15)

**B-J**, IPGTT (**B**) and ITT(**C**) tests, plasma triglyceride (**D**), total cholesterol (TC) (**E**), HDL-c (**F**) and LDL-c (**G**), representative images of H&E staining of iWAT, eWAT and liver (**H**), BAT, iWAT and eWAT content (**I**), total physical activities (**J**) and respiratory exchange ratio (RER) (**K**) of KI^cas9^ mice and WT littermates under HFD (**B**, **C**, n = 17:17; **D-G**, n = 10:10; **H**, n = 8:15; **I**, Scale bar, 100 μm; **J**, **K**, n = 8:8).

**L**, The validated diet-induced adiposity phenotype of KI^cas9^ mice (with F3 generation) under HFD challenge initiating from 11-week-old to 27-week-old (n = 18:19 for KI^cas9^ and WT, respectively).

**M**, Body weight curve of female KI^cas9^ and wild-type littermate mice under NCD (n = 21:20).

**N-Q**, Body weight curve (**N**), average food intake (**O**), body composition (**P**), and mass percentage of indicated fat tissue (**Q**) of female KI^cas9^ and wild-type littermates under HFD challenge initiating from 8-week-old to 20-week-old. (**N**, n = 9:8; **O**, n = 5:4, average food intake per mouse of individual cage; **P**, **Q**, n = 8:8).

Data are mean ± s.e.m. of biologically independent samples; unpaired two-sided Student’s *t*-test. * *P* < 0.05.

**Figure S3 | The homologous rs1421085 T>C variant protests HFD-induced obesity but shows no change in body weight under chow diet in KI^ES^ mice.**

**A**, Body weight of male KI^ES^ and WT mice aged 8- and 23-week-old under NCD (n = 13:10).

**B**, Body weight curve of female KI^ES^ and WT mice under NCD (n = 8:8).

**C-D**, BAT, iWAT and eWAT content (**C**), representative images of H&E staining of iWAT, eWAT and liver (**D**) of male KI^ES^ and wild-type littermates under HFD (**C**, n = 11:16; **D**, scale bar, 100 μm).

**E-K**, Body weight curve (**E**), average food intake (**F**), body composition (**G**) and mass percentage of indicated fat tissues (**H**), representative images of H&E staining of indicated tissues (**I**, **J**) and TEM of BAT (**K**) of female KI^ES^ and wild-type littermate mice under HFD challenge initiating from 10-week-old to 24-week-old (**E**, n = 16:13; **F**, n = 5:4 cages; **G, H,** n = 10–13; **I**, **J**, scale bar, 100 μm **K**, scale bar, 1 μm).

Data are mean ± s.e.m. of biologically independent samples; unpaired two-sided Student’s *t*-test. * *P* < 0.05

**Figure S4 | Brown adipocyte-specific knock-in homologous rs1421085 T>C variant enhances thermogenesisresists HFD-induced adiposity.**

**A-E**, The mRNA expression of *Fto* (**A**), *Irx3* (**B**), *Irx5* (**C**), *Irx6* (**D**), and *Rpgrip1l* (**E**) in the key tissues related to energy hemostasis of homologous rs1421085^fl/fl^ (KI^fl/fl^) and wild-type mice (4-5 weeks old male mice; n = 3:4).

**F**, **G**, Body weight curve (**F**) and mass percentage (**G**) of *Ucp1*-KI^fl/fl^ mice and KI^fl/fl^ littermates under NCD (n = 12:7).

**H**, The representative H&E images of iWAT, eWAT and liver in *Ucp1*-KI^fl/fl^ mice and KI^fl/fl^ littermates under HFD (Scale bar, 100 μm).

Data are mean ± s.e.m. of biologically independent samples.

**Figure S5 | The mRNA expression of potential downstream effectors of rs1421085 T>C variation.**

**A-D**, The mRNA expression of *Irx3*, *Irx5*, *Irx6*, and *Rpgrip1l* in BAT tissues and induced mature brown adipocytes of indicated models (KI^cas9^, n = 6:7; KI^ES^, n = 7:13; *Ucp1*-KI^fl/fl^, n = 15:7; induced brown adipocytes derived from KI^ES^ BAT SVF, n = 4:3).

Data are mean ± s.e.m. of biologically independent samples; unpaired two-sided Student’s *t*-test. * *P* < 0.05, ** *P* < 0.01.

**Figure S6 | FTO enhances thermogenic capacity of brown adipocytes and partially medicates the increased thermogenesis in rs1421085 T>C knockin model.**

**A**, The mRNA expression of *Fto* and thermogenesis-related genes at different time point during brown adipocyte differentiation derived from BAT SVFs (n = 5).

**B**, Correlation analysis of the mRNA levels of *Fto* and *Ucp1* in (**A**).

**C-E**, Oil Red O staining (**C**), OCR measurement (**D**, **E**) of induced brown adipocytes derived from BAT SVFs with *Fto* knockdown.

**F**, Oil Red O staining of induced brown adipocytes derived from BAT SVFs with *Fto* overexpression.

**G**, The m6A abundance in induced mature brown adipocytes in the presence of Entacapone (ENT, 50 μM) or vehicle (n = 3).

**H**, The mRNA expression of thermogenesis-related genes in induced mature brown adipocytes in the presence of ENT or vehicle (n = 4).

**I**, **J**, The protein levels (**I**) and quantification analysis (**J**) of FTO and UCP1 in induced mature brown adipocytes in the presence of ENT or vehicle (n = 3).

**C-F**, The SVFs were infected with lentivirus 2 days before induction; MOI = 50;

**C-G C**, **F**, Scale bar, 100 μm.

Data are mean ± s.e.m. of biologically independent samples; unpaired two-sided Student’s *t*-test. * *P* < 0.05, ** *P* < 0.01, *** *P* < 0.001.

**Figure S7 | *Fto* deficiency in BAT impairs thermogenesis and resists HFD-induced adiposity.**

**A**, The mRNA expression of *Fto* in indicated adipose tissues in 4-week-old female *Fto*^fl/fl^;*Myf5*-cre (MFKO) mice and *Fto*^fl/fl^ littermates (n = 4:4).

**B-D**, Body weight (**B**), body composition (**C**) and average food intake (**D**) of MFKO mice and littermate controls under HFD challenge initiating from 9-week-old to 19-week-old (n = 9:12).

**E-K**, Whole-day and 12h O_2_ consumption (**E, F**), CO_2_ production (**G, H**), heat generation (**I**), RER (**J**) and total physical activities (**K**) were collected in MFKO mice and littermate controls under HFD challenge (n = 8:8; **E-I**, normalized to the body weight^0.75^).

**L**, **M**, The protein expression (**L**) quantification analysis (**M**) and of FTO, UCP1 and PGC-1α in induced mature brown adipocytes derived from BAT SVFs of MFKO and littermate control mice (n = 4:4).

The data were presented as means ± s.e.m.; unpaired two-sided Student’s *t*-test. * *P* < 0.05, ** *P <* 0.01, *** *P* < 0.001.

**Figure S8 | FTO protein stabilizes Ucp1 mRNA and increase UCP1 protein expression.**

**A-D**, RIP-qPCR analysis of the interaction of FTO protein and target mRNA in induced mature brown adipocytes. Representative immunoblots showing the products of IP by FTO antibody, with IgG as a negative control (**A**); AGE (**B**) and qPCR analysis (**C, D**) of *Ucp1* and *C/ebpa* mRNA abundance of FTO-IP products (n = 4). The enrichment of *Ucp1* and *C/ebpa* mRNA were normalized to input (7.7%).

**E-F**, Half-life analysis of *Ucp1* (**E**) and *C/ebpa* (**F**) mRNA in induced mature brown adipocytes with *Fto* knockdown. The BAT SVFs were infected with lentivirus 2 days before induction (MOI = 75); Actinomycin D (5 μg/ml) was added on the eighth day of induction; total RNA was isolated at indicated time points (n = 3-4); Calculated half-times t_1/2_ = In2/K_decay_.

**G**-**H**, The protein expression (**G**) and quantification analysis (**H**) of FTO and UCP1 in induced mature brown adipocytes derived from BAT SVFs of KI^ES^ and wild-type littermate mice in the presence of ENT or vehicle (n = 3:3).

The data were presented as means ± s.e.m.; unpaired two-sided Student’s *t*-test. * *P* < 0.05, ** *P* < 0.01, *** *P* < 0.001.

**Figure S9 | Surgical excision of intrascapular BAT largely reverses HFD-resistance phenotype of KI^cas9^ mice.**

**A**, Representative anatomy images showed no obvious regeneration of intrascapular BAT 16 weeks after BAT-surgical deletion in KI^cas9^ (KI^cas9 ΔBAT^) and WT littermate (WT^ΔBAT^) mice. White dotted circle indicated interscapular BAT (iBAT) region.

**B-F**, Body weight curve (**B**), average food intake (**C**), body composition (**D**), mass percentage of indicated tissues (**E**), and representative H&E images of iWAT, eWAT and liver (**F**) in KI^cas9 ΔBAT^ mice and WT^ΔBAT^ littermates after HFD (Scale bar, 100 μm).

BAT-surgical deletion was performed to mice at 6-week-old, and HFD was provided to mice at 8-week-old; n = 7:9.

The data were presented as means ± s.e.m.; unpaired two-sided Student’s *t*-test. * *P* < 0.05, ** *P* < 0.01.

**Figure S10 | The association of human FTO rs1421085 T>C variation frequency with BMI.**

**A**, **B**, The Forest plot of the association between BMI and human FTO rs1421085 polymorphism in populations with ages older than 8 year-old (**A**) and younger than 8 year-old (**B**), respectively. Effect allele, rs1421085_C; reference allele, rs1421085_T.

**Figure S11 | The FTO and UCP1 were colocalized in human embryonic brown adipose tissue.** Representative immunofluorescence (IF) images of FTO (green) and UCP1 (red) in human embryonic brown adipose tissue from one 24-week-old donor (left) and one 20-week-old donor (right). The nucleus was stained with DAPI (blue), lipid droplets was stained with PLIN1 (pink). Top (PLIN1, UCP1, FTO, PLIN1/FTO/UCP1/PLIN1) scale bar, 200μm; bottom (PLIN1/FTO/UCP1/PLIN1) scale bar, 20μm.

**Figure S12 | The association of human FTO rs1421085 T>C variation frequency with ambient environmental temperature, elevation, and latitude.**

**A**, **B**, The correlation analysis of effect allele frequencies and CL-corrected (**A**) or CL-AVG corrected (**B**) annual mean earth skin temperature.

**C**, **D**, The correlation analysis of effect allele frequency and elevation (**C**) or longitude (**D**).

**A-D**, Colors and symbols represented populations of different continents. Details of populations are listed in **Table S5.**

**Figure S13 |** Sanger sequencing of the 6 kb DNA region centered on the homologous rs1421085 locus in WT and KI^cas9^ mice. The rs1421085 locus was emphasized in bold.

