## Supplementary material for "The rs1421085 variant within *FTO* promotes but not inhibits thermogenesis and is potentially associated with human migration": figure S 1-13

### Figure S1

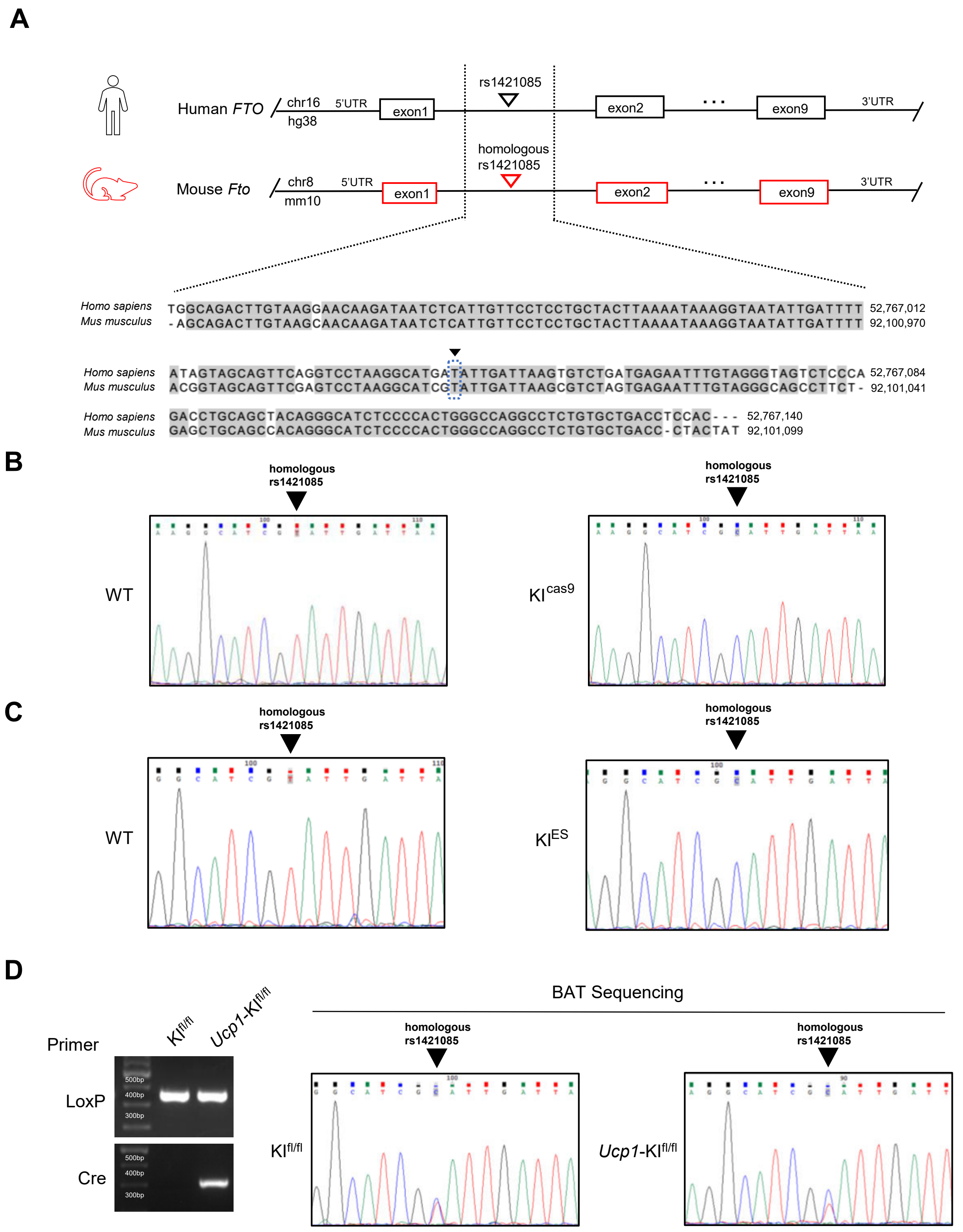
A
rs1421085
Human FTO
chr16
hg38
5'UTR
exon1
exon2
exon9
3'UTR
homologous
rs1421085
Mouse Fto
chr8
mm10
5'UTR
exon1
exon2
exon9
3'UTR
Homo sapiens TGGCAGACTTGTAAGGAACAAGATAATCTCATTGTTCCTCCTGCTACTTAAAATAAAGGTAATATTGATTTT 52,767,012
Mus musculus -AGCAGACTTGTAAGCAACAAGATAATCTCATTGTTCCTCCTGCTACTTAAAATAAAGGTAATATTGATTTT 92,100,970
Homo sapiens ATAGTAGCAGTTCAGGTCCTAAGGCATGATATTGATTAAGTGTCTGATGAGAATTTGTAGGGTAGTCTCCCA 52,767,084
Mus musculus ACGGTAGCAGTTCGAGTCCTAAGGCATCGTATTGATTAAGCGTCTAGTGAGAATTTGTAGGGCAGCCTTCT- 92,101,041
Homo sapiens GACCTGCAGCTACAGGGCATCTCCCCACTGGGCCAGGCCTCTGTGCTGACCTCCAC--- 52,767,140
Mus musculus GAGCTGCAGCCACAGGGCATCTCCCCACTGGGCCAGGCCTCTGTGCTGACC-CTACTAT 92,101,099
B
homologous rs1421085
WT
homologous rs1421085
KI cas9
C
homologous rs1421085
WT
homologous rs1421085
KI ES
D
Primer
KI fl/fl
Ucp1-KI fl/fl
LoxP
Cre
500bp
400bp
300bp
BAT Sequencing
homologous rs1421085
KI fl/fl
homologous rs1421085
Ucp1-KI fl/fl

### Figure S2

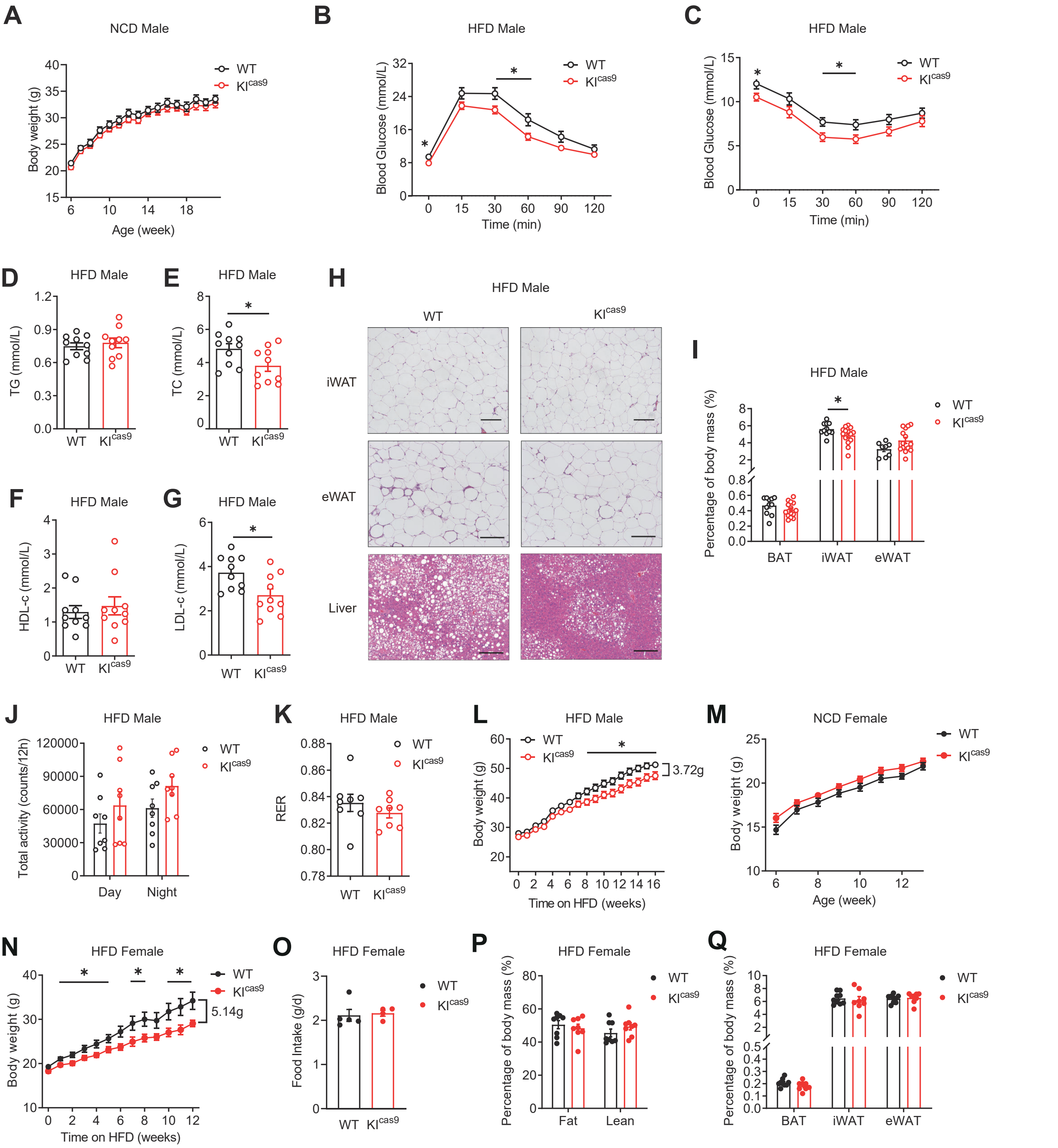
A
NCD Male
Body weight (g)
Age (week)
WT
KIcas9
B
HFD Male
Blood Glucose (mmol/L)
Time (min)
C
HFD Male
Blood Glucose (mmol/L)
Time (min)
D
HFD Male
TG (mmol/L)
E
HFD Male
TC (mmol/L)
F
HFD Male
HDL-c (mmol/L)
G
HFD Male
LDL-c (mmol/L)
H
HFD Male
iWAT
eWAT
Liver
I
HFD Male
Percentage of body mass (%)
BAT
iWAT
eWAT
J
HFD Male
Total activity (counts/12h)
Day
Night
K
HFD Male
RER
L
HFD Male
Body weight (g)
Time on HFD (weeks)
3.72g
M
NCD Female
Body weight (g)
Age (week)
N
HFD Female
Body weight (g)
Time on HFD (weeks)
5.14g
O
HFD Female
Food Intake (g/d)
P
HFD Female
Percentage of body mass (%)
Fat
Lean
Q
HFD Female
Percentage of body mass (%)
BAT
iWAT
eWAT

Figure S3

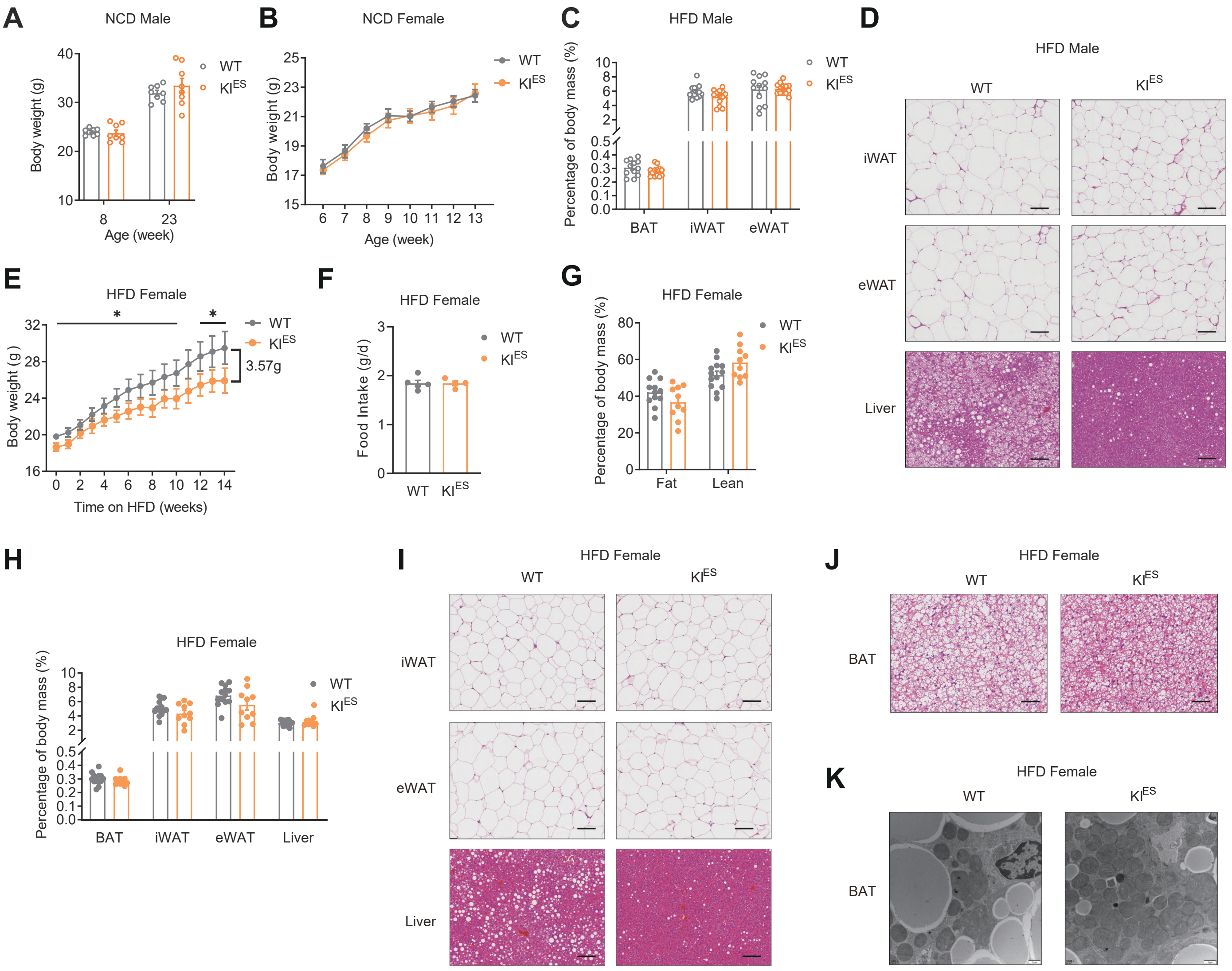

A
NCD Male
Body weight (g)
Age (week)
WT
KI^ES
B
NCD Female
Body weight (g)
Age (week)
C
HFD Male
Percentage of body mass (%)
BAT
iWAT
eWAT
D
HFD Male
iWAT
eWAT
Liver
E
HFD Female
Body weight (g)
Time on HFD (weeks)
3.57g
F
HFD Female
Food Intake (g/d)
G
HFD Female
Percentage of body mass (%)
Fat
Lean
H
HFD Female
BAT
iWAT
eWAT
Liver
I
HFD Female
J
HFD Female
BAT
K
HFD Female
BAT

Figure S4

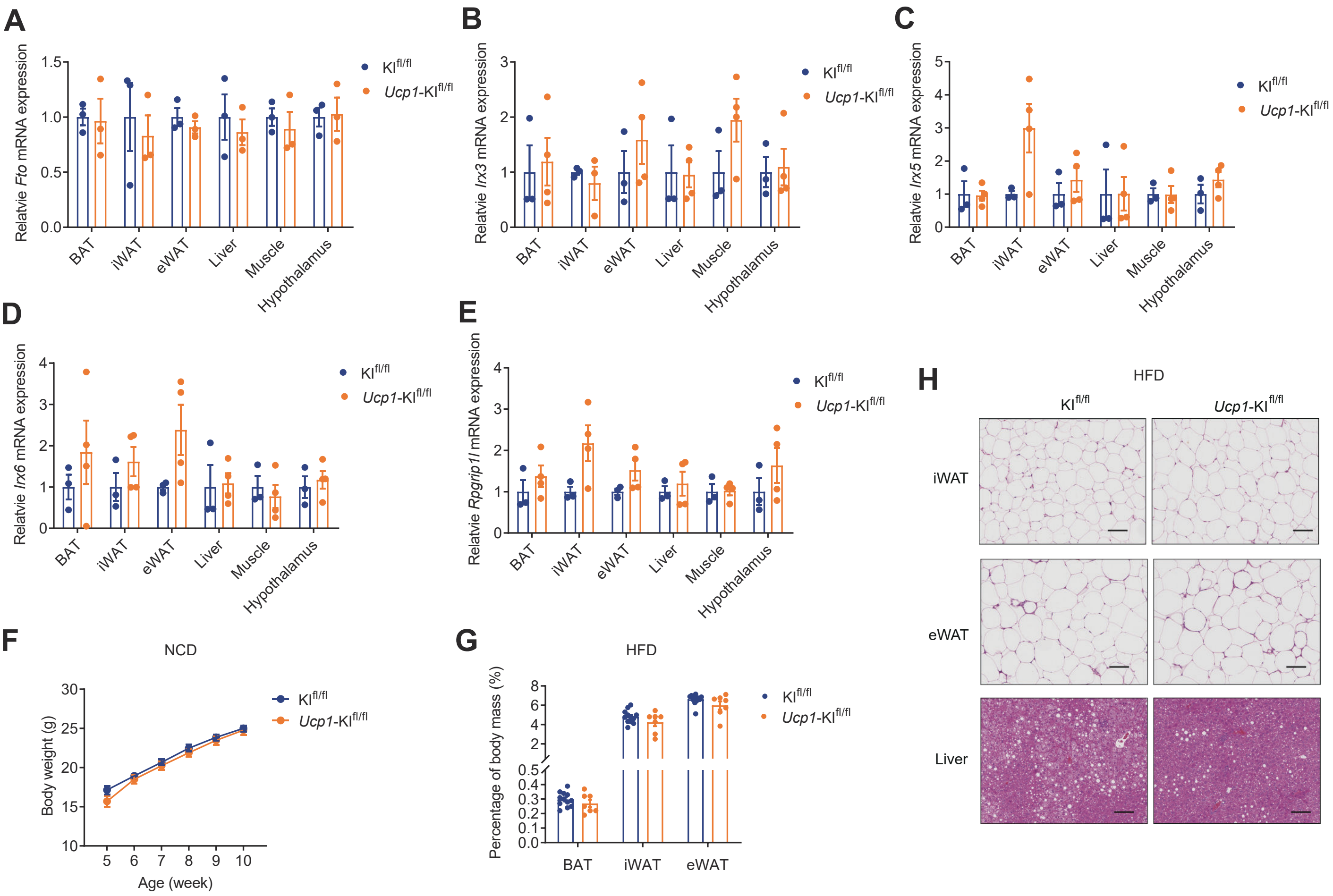

A
Relatvie Fto mRNA expression
Klfl/fl
Ucp1-Klfl/fl
BAT
iWAT
eWAT
Liver
Muscle
Hypothalamus
B
Relatvie Irx3 mRNA expression
C
Relatvie Irx5 mRNA expression
D
Relatvie Irx6 mRNA expression
E
Relatvie Rpgrip1l mRNA expression
F
NCD
Body weight (g)
Age (week)
G
HFD
Percentage of body mass (%)
H
HFD
iWAT
eWAT
Liver

### Figure S5

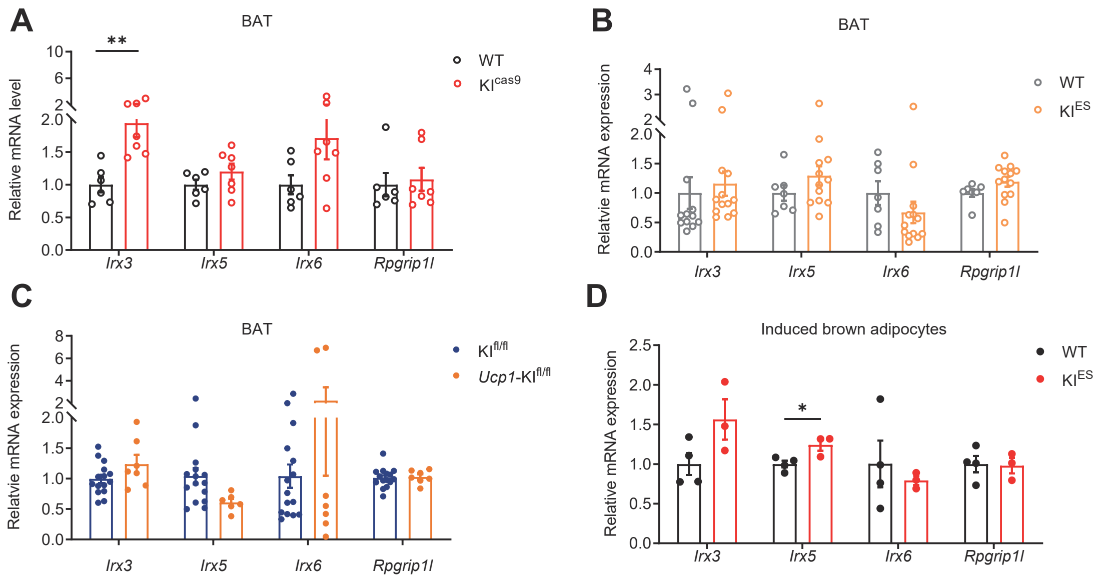

A
BAT
**
WT
KIcas9
Relative mRNA level
Irx3
Irx5
Irx6
Rpgrip1l
B
BAT
WT
KIES
Relatvie mRNA expression
C
BAT
KIfl/fl
Ucp1-KIfl/fl
Relatvie mRNA expression
D
Induced brown adipocytes
WT
KIES
*
Relative mRNA expression

### Figure S6

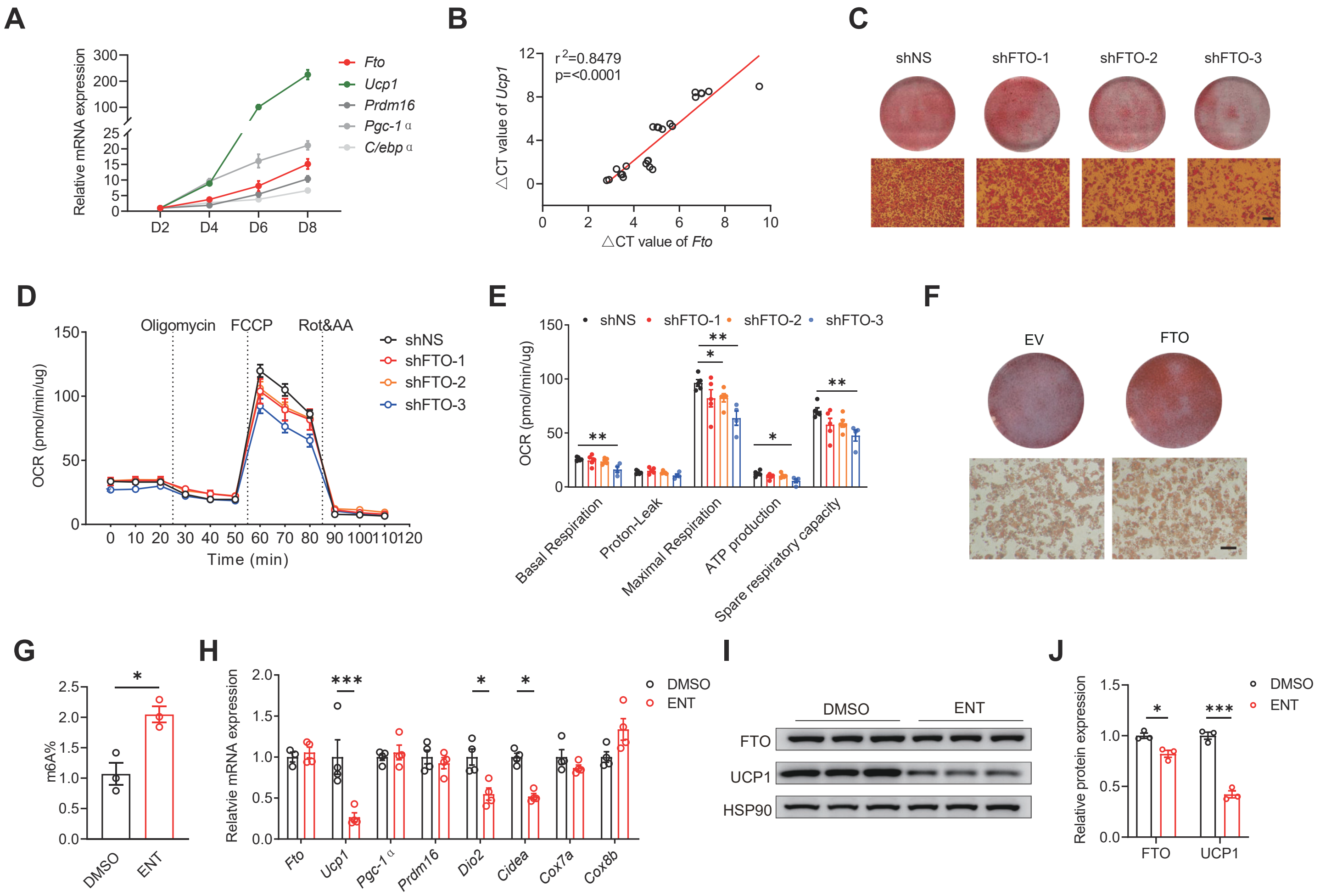

A
Relative mRNA expression
Fto
Ucp1
Prdm16
Pgc-1α
C/ebpα
D2
D4
D6
D8
B
r2=0.8479
p=<0.0001
△CT value of Ucp1
△CT value of Fto
C
shNS
shFTO-1
shFTO-2
shFTO-3
D
Oligomycin
FCCP
Rot&AA
OCR (pmol/min/ug)
Time (min)
shNS
shFTO-1
shFTO-2
shFTO-3
E
OCR (pmol/min/ug)
Basal Respiration
Proton-Leak
Maximal Respiration
ATP production
Spare respiratory capacity
F
EV
FTO
G
m6A%
DMSO
ENT
H
Relatvie mRNA expression
Fto
Ucp1
Pgc-1α
Prdm16
Dio2
Cidea
Cox7a
Cox8b
DMSO
ENT
I
DMSO
ENT
FTO
UCP1
HSP90
J
Relative protein expression
FTO
UCP1
DMSO
ENT

### Figure S7

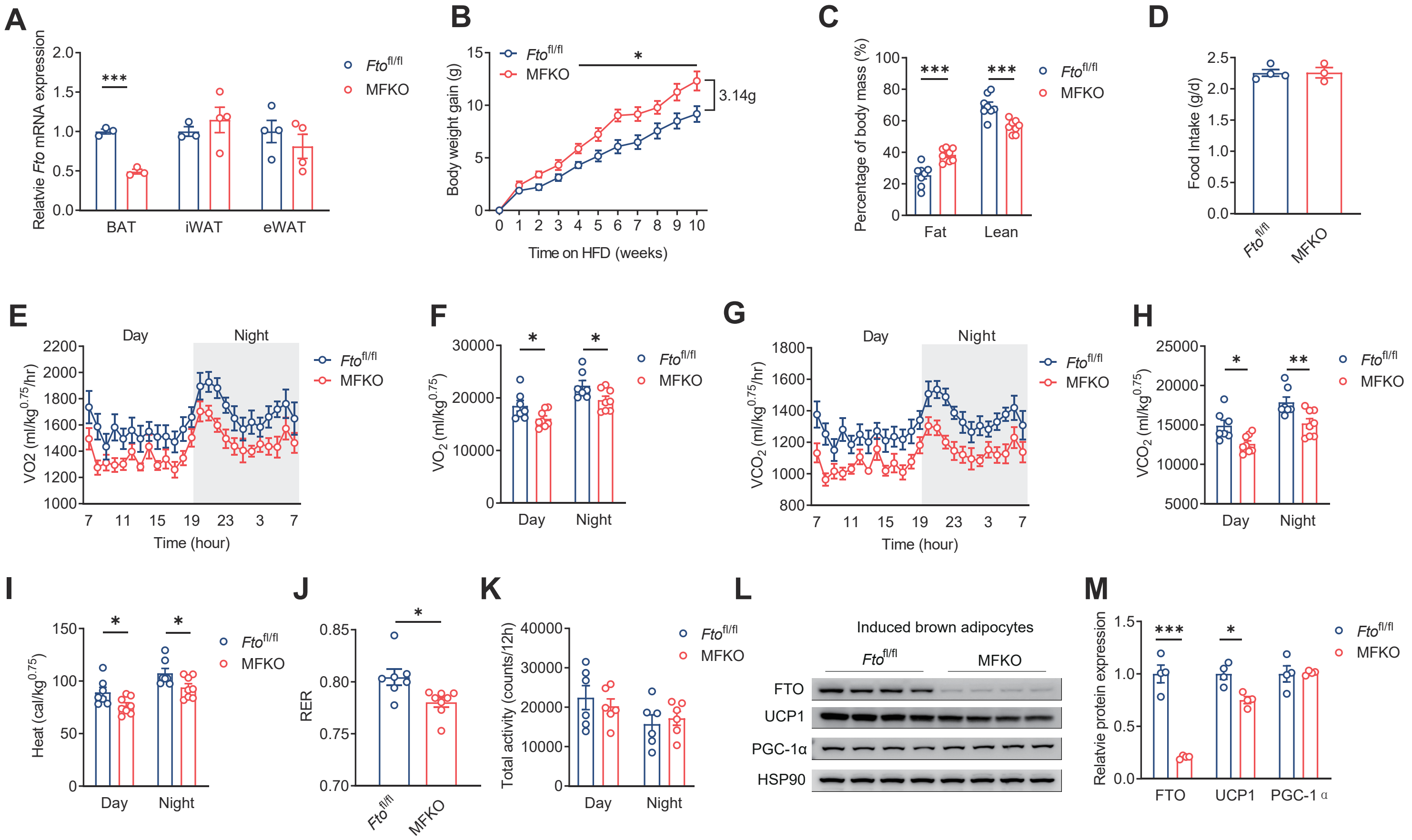
A
Relatvie Fto mRNA expression
BAT
iWAT
eWAT
Fto fl/fl
MFKO
B
Body weight gain (g)
Time on HFD (weeks)
3.14g
C
Percentage of body mass (%)
Fat
Lean
D
Food Intake (g/d)
E
VO2 (ml/kg0.75/hr)
Day
Night
Time (hour)
F
VO2 (ml/kg0.75)
G
VCO2 (ml/kg0.75/hr)
H
VCO2 (ml/kg0.75)
I
Heat (cal/kg0.75)
J
RER
K
Total activity (counts/12h)
L
Induced brown adipocytes
FTO
UCP1
PGC-1α
HSP90
M
Relatvie protein expression
FTO
UCP1
PGC-1α

Figure S8

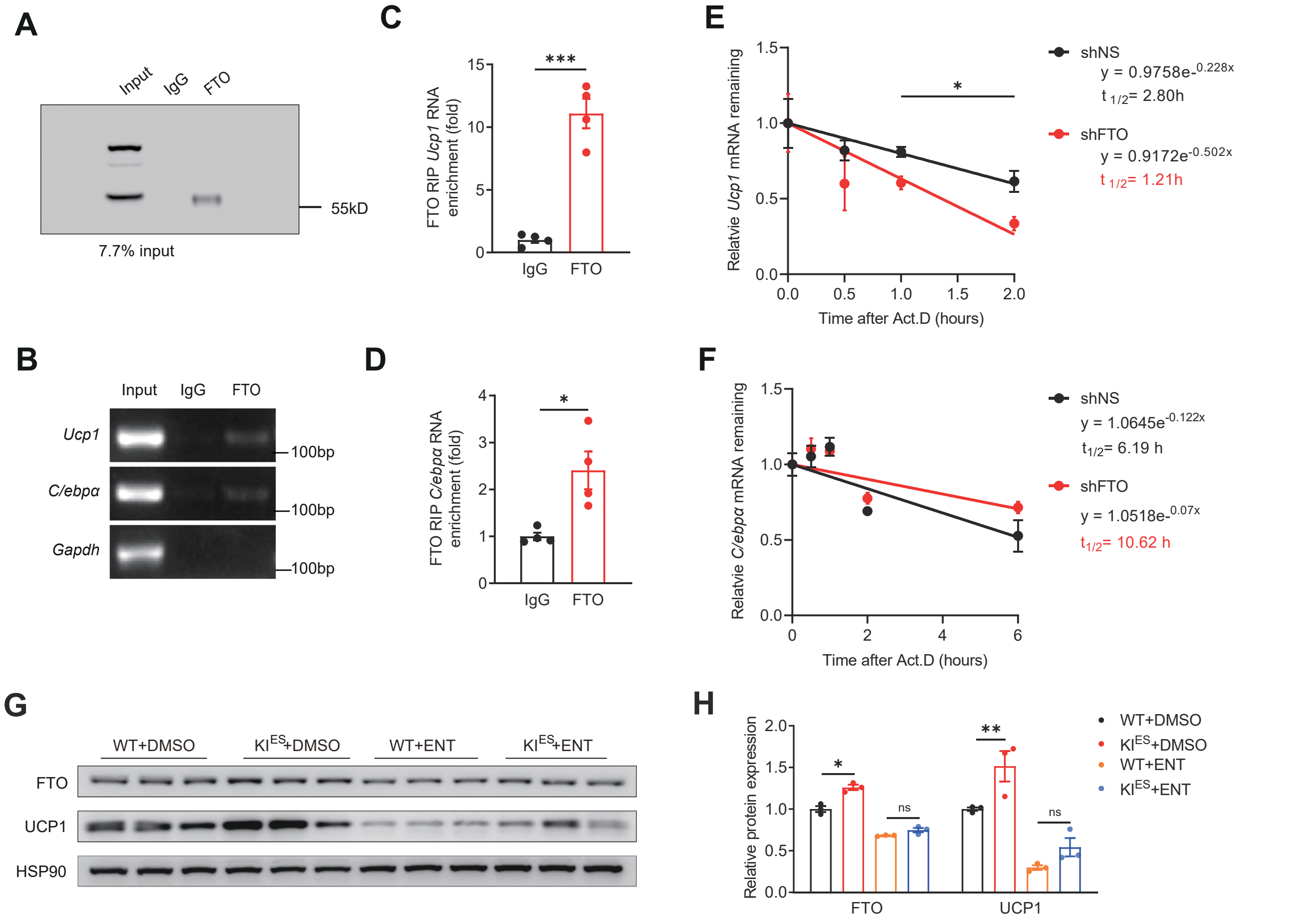

A
Input
IgG
FTO
55kD
7.7% input
B
Input
IgG
FTO
Ucp1
C/ebpα
Gapdh
100bp
100bp
100bp
C
FTO RIP Ucp1 RNA enrichment (fold)
***
IgG
FTO
D
FTO RIP C/ebpα RNA enrichment (fold)
*
IgG
FTO
E
Relatvie Ucp1 mRNA remaining
Time after Act.D (hours)
*
shNS
y = 0.9758e-0.228x
t 1/2= 2.80h
shFTO
y = 0.9172e-0.502x
t 1/2= 1.21h
F
Relatvie C/ebpα mRNA remaining
Time after Act.D (hours)
shNS
y = 1.0645e-0.122x
t1/2= 6.19 h
shFTO
y = 1.0518e-0.07x
t1/2= 10.62 h
G
WT+DMSO
KIES+DMSO
WT+ENT
KIES+ENT
FTO
UCP1
HSP90
H
Relative protein expression
FTO
UCP1
ns
ns
WT+DMSO
KIES+DMSO
WT+ENT
KIES+ENT

Figure S9

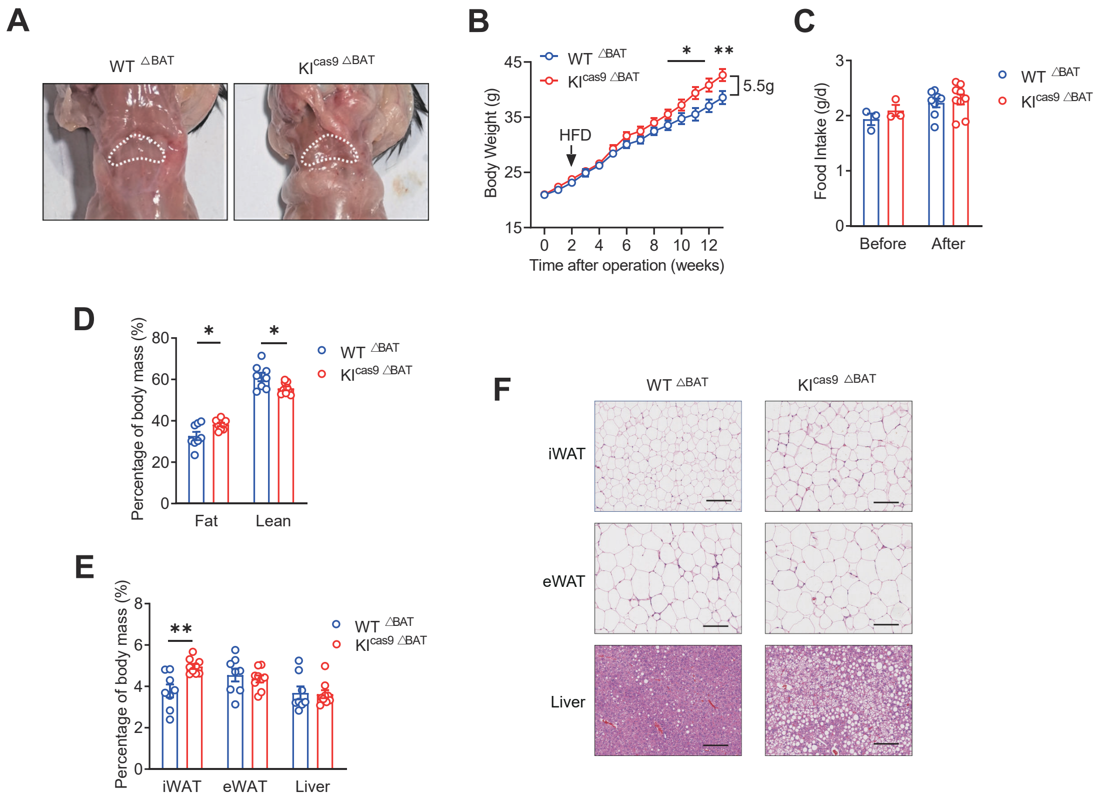

A
WT △BAT
KIcas9 △BAT
B
WT △BAT
KIcas9 △BAT
Body Weight (g)
HFD
5.5g
Time after operation (weeks)
C
Food Intake (g/d)
Before
After
D
Percentage of body mass (%)
Fat
Lean
E
Percentage of body mass (%)
iWAT
eWAT
Liver
F
WT △BAT
KIcas9 △BAT
iWAT
eWAT
Liver

Figure S10

**A**

| Study or Subgroup | Effect Allele Mean | Effect Allele SD | Effect Allele Total | Reference Allele Mean | Reference Allele SD | Reference Allele Total | Weight | Std. Mean Difference IV, Random, 95% CI |
|---|---|---|---|---|---|---|---|---|
| Albuquerque 2013 | 19.84 | 3.43 | 501 | 19 | 3.4 | 220 | 5.0% | 0.25 [0.09, 0.40] |
| Fang 2010 | 19.8 | 3.73 | 155 | 18.9 | 4.54 | 515 | 4.5% | 0.21 [0.03, 0.39] |
| González-Herrera 2019 | 20.15 | 4.41 | 211 | 19.3 | 4.11 | 298 | 4.6% | 0.20 [0.02, 0.38] |
| Hakanen 2009 | 20.54 | 3.34 | 291 | 20.3 | 3.65 | 148 | 4.1% | 0.07 [−0.13, 0.27] |
| Jacobsson 2008 | 27.82 | 1.04 | 653 | 27.6 | 0.5 | 307 | 5.7% | 0.24 [0.11, 0.38] |
| Labayen 2016 | 21.35 | 3.64 | 406 | 20.7 | 3.2 | 246 | 5.0% | 0.19 [0.03, 0.35] |
| Liliane 2019 | 23.15 | 4.25 | 457 | 23.47 | 4.4 | 299 | 5.4% | −0.07 [−0.22, 0.07] |
| Liu 2010 | 22.95 | 4.89 | 1376 | 22.5 | 4.37 | 599 | 6.8% | 0.09 [−0.00, 0.19] |
| Mangge 2011 | 28.08 | 7.14 | 265 | 26.5 | 7 | 106 | 3.5% | 0.22 [−0.00, 0.45] |
| Olza 2013 | 23.34 | 6.08 | 369 | 21.69 | 6.11 | 162 | 4.4% | 0.27 [0.09, 0.46] |
| Pausova 2009 | 21.54 | 3.17 | 273 | 20.7 | 2.91 | 212 | 4.5% | 0.27 [0.09, 0.45] |
| Rivas 2018 | 23.01 | 4.86 | 111 | 23.1 | 4.8 | 147 | 3.2% | −0.02 [−0.27, 0.23] |
| Rutters 2011 | 19.6 | 1.9 | 56 | 18.3 | 2.9 | 45 | 1.6% | 0.54 [0.14, 0.94] |
| Scott 2010 | 15.04 | 6.96 | 772 | 16.3 | 6.77 | 176 | 4.9% | −0.18 [−0.35, −0.02] |
| Shinozaki 2018 | 19.35 | 2.7 | 52 | 18.9 | 2.7 | 104 | 2.1% | 0.17 [−0.17, 0.50] |
| Tanofsky-Kraff 2009 | 25.87 | 6.26 | 109 | 22.85 | 7.96 | 99 | 2.7% | 0.42 [0.15, 0.70] |
| Vasan 2013 | 20.68 | 4.1 | 636 | 20.7 | 3.9 | 580 | 6.4% | −0.00 [−0.12, 0.11] |
| Wardle 2008 | 17.81 | 7.53 | 2128 | 17.18 | 7.63 | 1209 | 7.6% | 0.08 [0.01, 0.15] |
| Wu 2014 | 22.58 | 3.89 | 86 | 21.87 | 3.68 | 310 | 3.3% | 0.19 [−0.05, 0.43] |
| Xi 2010 | 22.54 | 2.89 | 785 | 21.7 | 5.21 | 2718 | 7.3% | 0.18 [0.10, 0.25] |
| Yang 2014 | 21.43 | 5.88 | 942 | 20.2 | 5.6 | 2982 | 7.5% | 0.22 [0.14, 0.29] |
| **Total (95% CI)** | | | **10634** | | | **11482** | **100.0%** | **0.15 [0.09, 0.20]** |

Heterogeneity: $Tau^2 = 0.01; Chi^2 = 57.64, df = 20 (P < 0.0001); I^2 = 65\%$
Test for overall effect: Z = 5.21 (P < 0.00001)

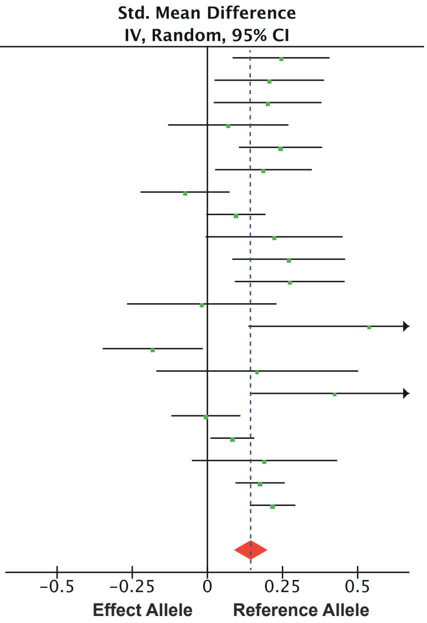


**B**

| Study or Subgroup | Effect Allele Mean | Effect Allele SD | Effect Allele Total | Reference Allele Mean | Reference Allele SD | Reference Allele Total | Weight | Std. Mean Difference IV, Random, 95% CI |
|---|---|---|---|---|---|---|---|---|
| Frayling 2008 | 16.84 | 1.93 | 3807 | 16 | 1.78 | 2162 | 26.6% | 0.45 [0.39, 0.50] |
| Lauria 2012 | 16.47 | 2.22 | 2824 | 16.18 | 2.33 | 1581 | 26.5% | 0.13 [0.07, 0.19] |
| Mook-Kanamori 2011 | 16.7 | 1.3 | 420 | 16.8 | 1.3 | 283 | 24.4% | −0.08 [−0.23, 0.07] |
| Zhang 2014 | 18.78 | 4.25 | 117 | 18.7 | 4 | 440 | 22.6% | 0.02 [−0.18, 0.22] |
| **Total (95% CI)** | | | **7168** | | | **4466** | **100.0%** | **0.14 [−0.10, 0.38]** |

Heterogeneity: $Tau^2 = 0.06; Chi^2 = 88.79, df = 3 (P < 0.00001); I^2 = 97\%$
Test for overall effect: Z = 1.13 (P = 0.26)

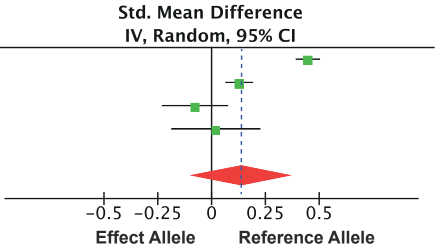

Figure S13

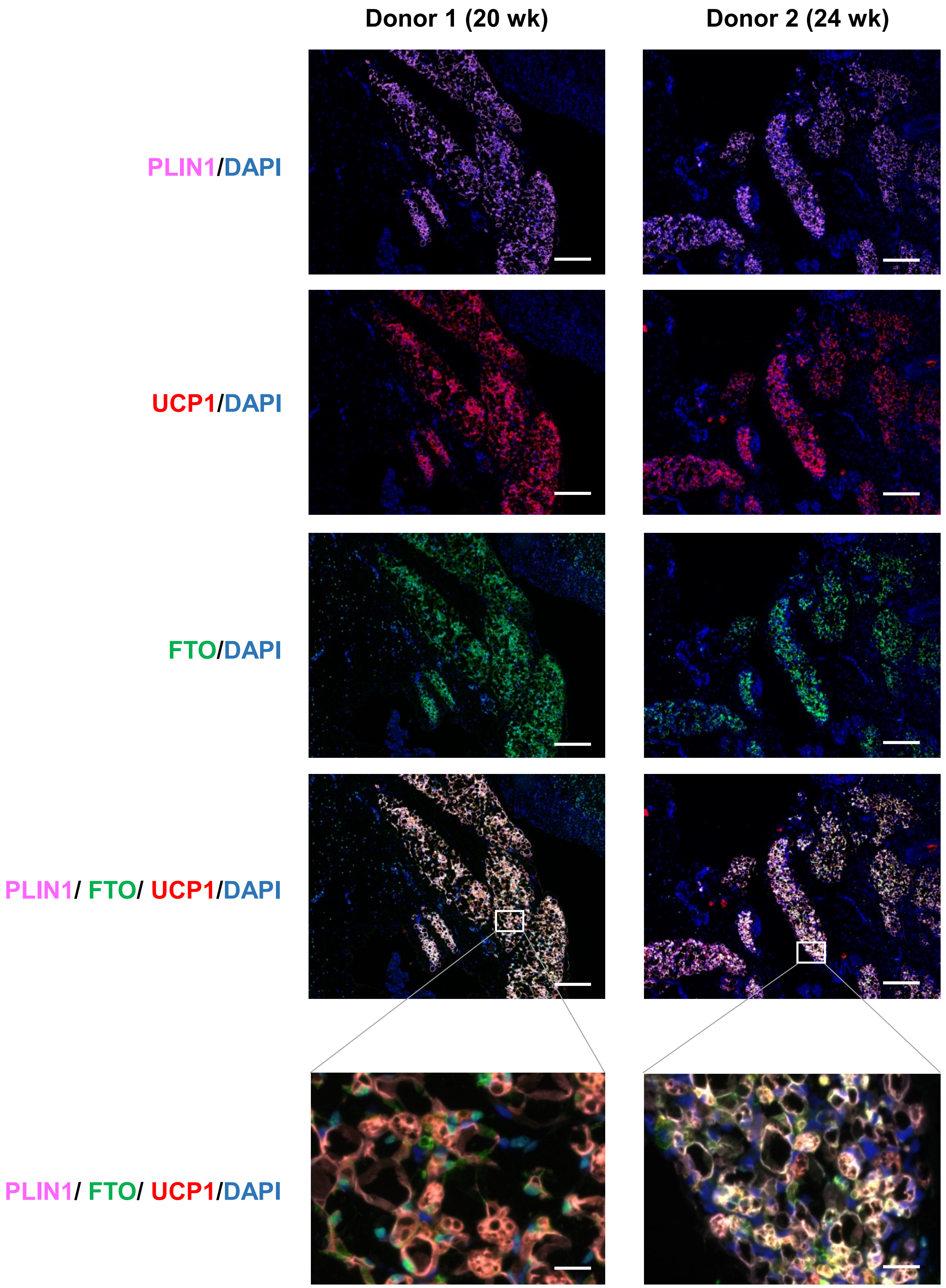

Donor 1 (20 wk)
Donor 2 (24 wk)
PLIN1/DAPI
UCP1/DAPI
FTO/DAPI
PLIN1/ FTO/ UCP1/DAPI
PLIN1/ FTO/ UCP1/DAPI

Figure S12

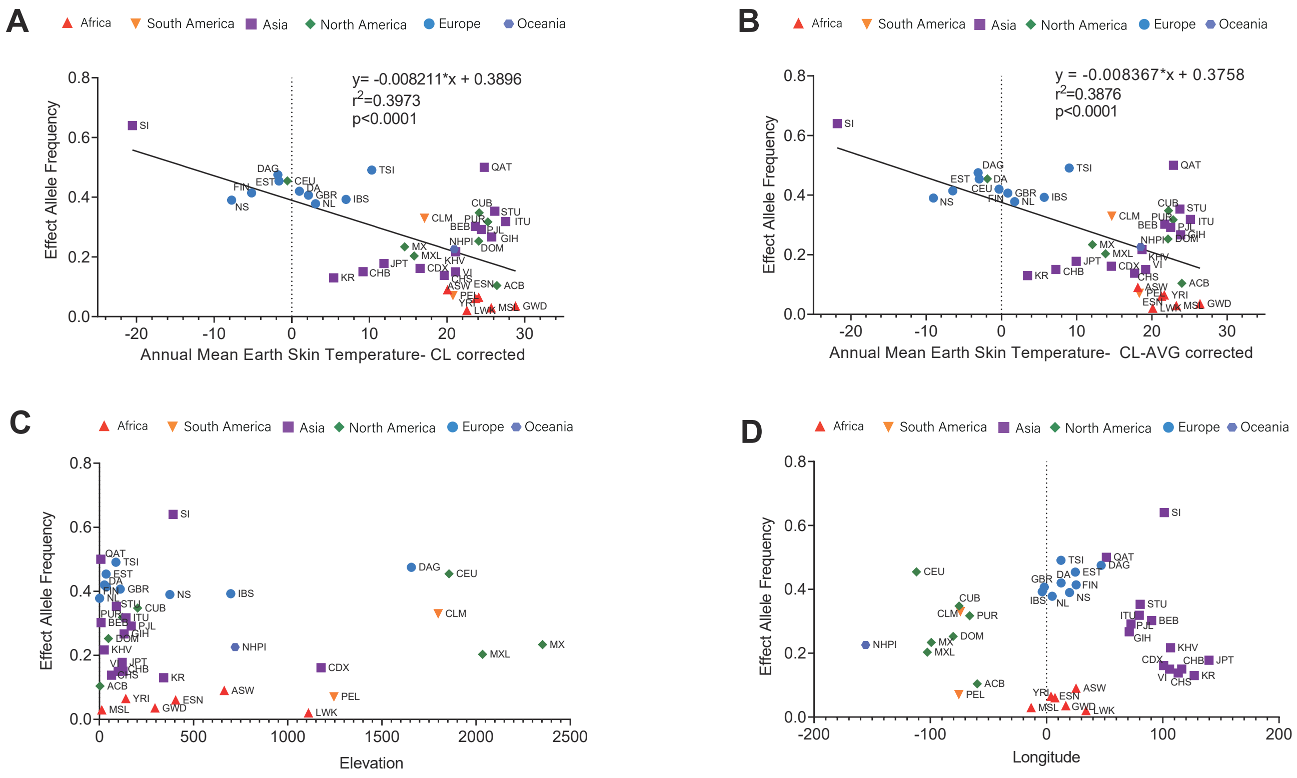
A
Africa
South America
Asia
North America
Europe
Oceania
Effect Allele Frequency
y= -0.008211*x + 0.3896
r2=0.3973
p<0.0001
Annual Mean Earth Skin Temperature- CL corrected
B
y = -0.008367*x + 0.3758
r2=0.3876
p<0.0001
Annual Mean Earth Skin Temperature- CL-AVG corrected
C
Elevation
D
Longitude

### Figure S13

WT 91371134 GAGGATGGGCTTTATCGTTTTAGGCTGAACCTGACACCGTATAACTCACAAGCCTGTGATCTTAAAAAACAAACAGAAAGGAAGTAGGAAAAAAGATGGAGAGGAAGCAGGAAAGGAGGGAGGGCGGAATGAAAGAAGGAAGGACAGAA 91371283
KI$^{cas9}$ 91371134 GAGGATGGGCTTTATCGTTTTAGGCTGAACCTGACACCGTATAACTCACAAGCCTGTGATCTTAAAAAACAAACAGAAAGGAAGTAGGAAAAAAGATGGAGAGGAAGCAGGAAAGGAGGGAGGGCGGAATGAAAGAAGGAAGGACAGAA 91371283

WT 91371284 TCTTTGCCTGGTGGCATTTCGTTGTTTGCTGTAGCTTATCACAAAAGAGGAGTGCCTTCTGGAGACAGTGACCTCCCTAAGAGCTGAGTTCTCACTGGTCCCGAGAAAAGGGCCTATAGCATCAGAATGATTGCTCTTGGTGGTACAAGT 91371433
KI$^{cas9}$ 91371284 TCTTTGCCTGGTGGCATTTCGTTGTTTGCTGTAGCTTATCACAAAAGAGGAGTGCCTTCTGGAGACAGTGACCTCCCTAAGAGCTGAGTTCTCACTGGTCCCGAGAAAAGGGCCTATAGCATCAGAATGATTGCTCTTGGTGGTACAAGT 91371433

WT 91371434 TTCAAAGTTTTAACTTAATATGTCATTAATGCACCTTAAAAGTTGTCACCTTATCTGTGAAAACAGAGGGCCGATTAGCAGAGAAGTGATCAGTATTGTTTTCAAGCAGAGTATGAAATGAATTTATTTCTTCACAGAATGCTATTTGAA 91371583
KI$^{cas9}$ 91371434 TTCAAAGTTTTAACTTAATATGTCATTAATGCACCTTAAAAGTTGTCACCTTATCTGTGAAAACAGAGGGCCGATTAGCAGAGAAGTGATCAGTATTGTTTTCAAGCAGAGTATGAAATGAATTTATTTCTTCACAGAATGCTATTTGAA 91371583

WT 91371584 ATGGGTTTGTGATGAATATCCTTAATAATGTTTAGTGAGAAAATAGAGCAATAAAGAAAAATATCTGGAACAATTGGAAGGTCTAGAAGATATTTATCAGATGATTTCTGATGTGTTCCAGTATAGAGAAGAACTATGACAGCTGGGCAT 91371733
KI$^{cas9}$ 91371584 ATGGGTTTGTGATGAATATCCTTAATAATGTTTAGTGAGAAAATAGAGCAATAAAGAAAAATATCTGGAACAATTGGAAGGTCTAGAAGATATTTATCAGATGATTTCTGATGTGTTCCAGTATAGAGAAGAACTATGACAGCTGGGCAT 91371733

WT 91371734 TGGGGGCTCGGGGTGGGGGTGGGGGAACATCTATGACCCCAGCACTCAGGTGGAGGAAGATGTTCAAGGCCAGCCTGGATTACAAACCAAGATCCTGTCTACCCCTTAAATAAAAGGACTAAAATAAGCATGCTAGCCCAATTACTTTTG 91371883
KI$^{cas9}$ 91371734 TGGGGGCTCGGGGTGGGGGTGGGGGAACATCTATGACCCCAGCACTCAGGTGGAGGAAGATGTTCAAGGCCAGCCTGGATTACAAACCAAGATCCTGTCTACCCCTTAAATAAAAGGACTAAAATAAGCATGCTAGCCCAATTACTTTTG 91371883

WT 91371884 GCCTTTTGCCCAATATTTTGAATGCAATTTTTTTCTTGTAGTCACAGACCTTTTAAGATGTCATGATGTCATGTGACTTATTAAAAATAATTATTGTAGGTGAGTAAAGATTGACAAGAGAAAAATTTGCTTCATTAAATTTTAATTTTA 91372033
KI$^{cas9}$ 91371884 GCCTTTTGCCCAATATTTTGAATGCAATTTTTTTCTTGTAGTCACAGACCTTTTAAGATGTCATGATGTCATGTGACTTATTAAAAATAATTATTGTAGGTGAGTAAAGATTGACAAGAGAAAAATTTGCTTCATTAAATTTTAATTTTA 91372033

WT 91372034 CTCTTGATTTATTTAGAAAGCTCCCAGATGAGAGGGTATTTTAACATAGTGATTCTATTTTTGTTTCTTGGGTCTTGTATTTTTAAGGGAGTTCCGATTGGCTGCAGCCATTGGCTTCTGAGGATCATTTGGCTCTGGGCCCCACTGTGA 91372183
KI$^{cas9}$ 91372034 CTCTTGATTTATTTAGAAAGCTCCCAGATGAGAGGGTATTTTAACATAGTGATTCTATTTTTGTTTCTTGGGTCTTGTATTTTTAAGGGAGTTCCGATTGGCTGCAGCCATTGGCTTCTGAGGATCATTTGGCTCTGGGCCCCACTGTGA 91372183

WT 91372184 GCACCCGGGAGTCTGACATGTGTTATGCTGAGCTACGTTATTTGTCCTTTGATAACACTGACCAAGTTGACAAGCTCCTGACATTTGTCCGGGTGAGGTAGAAATACTAGGAAGGAAAATTTAGTACTGCTCTCCTGTACAACTCAACTA 91372333
KI$^{cas9}$ 91372184 GCACCCGGGAGTCTGACATGTGTTATGCTGAGCTACGTTATTTGTCCTTTGATAACACTGACCAAGTTGACAAGCTCCTGACATTTGTCCGGGTGAGGTAGAAATACTAGGAAGGAAAATTTAGTACTGCTCTCCTGTACAACTCAACTA 91372333

WT 91372334 TTTTACCTGTGTGTATGATAGGAAAGGGAAATTTTACAGGGATTTTTGTATTTGTTCTGTAGCAATCCAGTTCACACAGAGGTTGCCATGTAGGGGCCCAGTCAGGAGAAAGTGGTGAAGCACTGTCTGGAACAGGAGACATCCAATTTA 91372483
KI$^{cas9}$ 91372334 TTTTACCTGTGTGTATGATAGGAAAGGGAAATTTTACAGGGATTTTTGTATTTGTTCTGTAGCAATCCAGTTCACACAGAGGTTGCCATGTAGGGGCCCAGTCAGGAGAAAGTGGTGAAGCACTGTCTGGAACAGGAGACATCCAATTTA 91372483

WT 91372484 ATCAGCAAGAGCAAGATTTTCTTTGTGTGTTCATTATTTTTGTTGCTAAGATGATTCCTTAGAGAGAGTCGGGACTTTAGAGTGTGAATGATGTCAGTTTCATCTCTGCTGCTTCCTTGTTTGATTTCTGCAAAGTCTTGGACCTAATAT 91372633
KI$^{cas9}$ 91372484 ATCAGCAAGAGCAAGATTTTCTTTGTGTGTTCATTATTTTTGTTGCTAAGATGATTCCTTAGAGAGAGTCGGGACTTTAGAGTGTGAATGATGTCAGTTTCATCTCTGCTGCTTCCTTGTTTGATTTCTGCAAAGTCTTGGACCTAATAT 91372633

WT 91372634 GCCAGTTTCTTACCTAGAAATGGGGCTTATTTGAATTTACCCCTGAGGCTCGATCAAACTTCAGAGTACTGACCATCATGCCCAGCTTCTGAGGAGAGGCTAACTCCTTACCCAGCTGTGTACCTGGCAGGTGGGTGGGCACAACGCCTT 91372783
KI$^{cas9}$ 91372634 GCCAGTTTCTTACCTAGAAATGGGGCTTATTTGAATTTACCCCTGAGGCTCGATCAAACTTCAGAGTACTGACCATCATGCCCAGCTTCTGAGGAGAGGCTAACTCCTTACCCAGCTGTGTACCTGGCAGGTGGGTGGGCACAACGCCTT 91372783

WT 91372784 ACCCAGAACACACGAGTGAGTGGACATTGGCCCCTCACCCAGCTCCTGGGTGGATCCTGTGCCTTTCCTGTTCATTTTCCTTTGCCTTGGAGCCAAACAGGCTTTAGCTAGGAGGTGAGCCTGCTCTTGATTGGTAGAGGGGATGAATTC 91372933
KI$^{cas9}$ 91372784 ACCCAGAACACACGAGTGAGTGGACATTGGCCCCTCACCCAGCTCCTGGGTGGATCCTGTGCCTTTCCTGTTCATTTTCCTTTGCCTTGGAGCCAAACAGGCTTTAGCTAGGAGGTGAGCCTGCTCTTGATTGGTAGAGGGGATGAATTC 91372933

WT 91372934 AGAAAGGCACAGAGTACTGTGGGCAGTCTTGTGAGGAGCTGAGGAAGGCTGTGTGTGTGTGTGTGTGTGTGTGTGTGTGTGTGTGTGTGTGTGAGAGAGAGAGAGAGAGAGAGATGGAGAGTGTGCAGGCAAGACGGGCCAAATCCACTCATTG 91373083
KI$^{cas9}$ 91372934 AGAAAGGCACAGAGTACTGTGGGCAGTCTTGTGAGGAGCTGAGGAAGGCTGTGTGTGTGTGTGTGTGTGTGTGTGTGTGTGTGTGTGTGTGTGAGAGAGAGAGAGAGAGAGAGATGGAGAGTGTGCAGGCAAGACGGGCCAAATCCACTCATTG 91373083

WT 91373084 GAGGCAGATTTGGTGATGGTGTCTGGAGACCCCATCTGAACCTCTGCTGTCAACACCGGCTTCATAGCATTGAAATCCAGGTCGAAGGAGCTGGGACTTGTTTTCTTCTTGACCTCTTATTGAAAATATTACTCTTTTGGAATGTTAAAA 91373233
KI$^{cas9}$ 91373084 GAGGCAGATTTGGTGATGGTGTCTGGAGACCCCATCTGAACCTCTGCTGTCAACACCGGCTTCATAGCATTGAAATCCAGGTCGAAGGAGCTGGGACTTGTTTTCTTCTTGACCTCTTATTGAAAATATTACTCTTTTGGAATGTTAAAA 91373233

WT 91373234 AGATTAAACCAATTTAGAAAAAAACCAAAGGTCTTTCTGTGCTCAGCTTCTCTCTCTGACTGAAGCGCTTTGCCAGTATGTTCAGCGTGACGGATATCAATACCAGATTTCCATGGTTACCCATGTTTTCCTTACAAGGAATCTAGCAGA 91373233
KI$^{cas9}$ 91373234 AGATTAAACCAATTTAGAAAAAAACCAAAGGTCTTTCTGTGCTCAGCTTCTCTCTCTGACTGAAGCGCTTTGCCAGTATGTTCAGCGTGACGGATATCAATACCAGATTTCCATGGTTACCCATGTTTTCCTTACAAGGAATCTAGCAGA 91373233

WT 91373233 GTATGTGCTAGTAGGTGCTCACTCTGTTCTTGGTGGTTTCTGTGGTAACAACACAGACCAGAACTGTTCCCCTCTAGGCTAACATTCTGTTAGAGGCAAGCCAGGAAGCAGGTAAAGAAGTGATGAGGAAGATGGCAGGAGGGGCTTTAG 91373533
KI$^{cas9}$ 91373233 GTATGTGCTAGTAGGTGCTCACTCTGTTCTTGGTGGTTTCTGTGGTAACAACACAGACCAGAACTGTTCCCCTCTAGGCTAACATTCTGTTAGAGGCAAGCCAGGAAGCAGGTAAAGAAGTGATGAGGAAGATGGCAGGAGGGGCTTTAG 91373533

WT 91373534 TTAGAGTGATCCGAAATAGCATCTTATATGGAGCCCCAAATACGACAACACGGGCTGAGGATGCTTGGATTTTTCATGTCCTGATAAGTCTTTTCATATAGGCGATGAAGGTGCTCTGTGGCCAGAATTTAAGGTCTTAGAAGCACCTTT 91373683
KI$^{cas9}$ 91373534 TTAGAGTGATCCGAAATAGCATCTTATATGGAGCCCCAAATACGACAACACGGGCTGAGGATGCTTGGATTTTTCATGTCCTGATAAGTCTTTTCATATAGGCGATGAAGGTGCTCTGTGGCCAGAATTTAAGGTCTTAGAAGCACCTTT 91373683

WT 91373684 AGTCTTGGGTAAATATTTAGATTCTATTACAACTCAATGGAGACCACAGTCAGGGTTAAGGTGAAGAATAACTGCTGATTTGTTATTTTATAAATATATCTGAGGGTGAGAATAGAATTAGGGAGGAAGTCTTGCCAAGGGTGCATTGAT 91373833
KI$^{cas9}$ 91373684 AGTCTTGGGTAAATATTTAGATTCTATTACAACTCAATGGAGACCACAGTCAGGGTTAAGGTGAAGAATAACTGCTGATTTGTTATTTTATAAATATATCTGAGGGTGAGAATAGAATTAGGGAGGAAGTCTTGCCAAGGGTGCATTGAT 91373833

WT 91373834 GTCTGGGACAGGAGGTGATTACAGAGGCCCATAGAAGGGGAGAGACTTAGGATCTAATCCTGAGGCAAAGTTGATGGGATTTCTTGTGAGACTGGATGAAGATGATTCAGAGACGAGAGGGACCAGAATGGCATCAGGGACTAGGTTGTG 91373983
KI$^{cas9}$ 91373834 GTCTGGGACAGGAGGTGATTACAGAGGCCCATAGAAGGGGAGAGACTTAGGATCTAATCCTGAGGCAAAGTTGATGGGATTTCTTGTGAGACTGGATGAAGATGATTCAGAGACGAGAGGGACCAGAATGGCATCAGGGACTAGGTTGTG 91373983

WT 91373984 TCTGTTGATGGCAATCAGAAGACTGAGGAAGAACAGTCCTTTGGTCTTCAGGTCCCCATGTGGCATCCAAGGGGCTGCTGGGAGGCAGGAATCTGAGAGCAGGAGGTCCAGTGTGCAGTTGCTGGATGGAAGCTTTGGGGCATCTTTCTT 91374133
KI$^{cas9}$ 91373984 TCTGTTGATGGCAATCAGAAGACTGAGGAAGAACAGTCCTTTGGTCTTCAGGTCCCCATGTGGCATCCAAGGGGCTGCTGGGAGGCAGGAATCTGAGAGCAGGAGGTCCAGTGTGCAGTTGCTGGATGGAAGCTTTGGGGCATCTTTCTT 91374133

WT 91374134 TAGCTTTCCTGAGCTAGTGTGTGTACCCTCAACTCTCCCTGTCTGTGGCCCTGCTGCCAGGCTGCCTCCGAGCCCAGCAAACTCATTCCTCCCAGGCAAAAGCAAAAGGTGACATACACCAGGAGCCAGATTGTCCACAGCAGACTTGTA 91374283
KI$^{cas9}$ 91374134 TAGCTTTCCTGAGCTAGTGTGTGTACCCTCAACTCTCCCTGTCTGTGGCCCTGCTGCCAGGCTGCCTCCGAGCCCAGCAAACTCATTCCTCCCAGGCAAAAGCAAAAGGTGACATACACCAGGAGCCAGATTGTCCACAGCAGACTTGTA 91374283

WT 91374284 AGCAACAAGATAATCTCATTGTTCCTCCTGCTACTTAAAATAAAGGTAATATTGATTTTACGGTAGCAGTTCGAGTCCTAAGGCATCG**T**ATTGATTAAGCGTCTAGTGAGAATTTGTAGGGCAGCCTTCTGAGCTGCAGCCACAGGGCAT 91374433
KI$^{cas9}$ 91374284 AGCAACAAGATAATCTCATTGTTCCTCCTGCTACTTAAAATAAAGGTAATATTGATTTTACGGTAGCAGTTCGAGTCCTAAGGCATCG**C**ATTGATTAAGCGTCTAGTGAGAATTTGTAGGGCAGCCTTCTGAGCTGCAGCCACAGGGCAT 91374433

WT 91374434 CTCCCCACTGGGCCAGGCCTCTGTGCTGACCCTACTATTAGAAGTGTTTAGGCTCTGGTCAGAACATTTATTTTACCTGATGGATGGCTTAATCCAGCCCGTCACCTTAATCTGACCTTAAAGATGGCAGTCCAACTCTTTTGTTCTGAG 91374583
KI$^{cas9}$ 91374434 CTCCCCACTGGGCCAGGCCTCTGTGCTGACCCTACTATTAGAAGTGTTTAGGCTCTGGTCAGAACATTTATTTTACCTGATGGATGGCTTAATCCAGCCCGTCACCTTAATCTGACCTTAAAGATGGCAGTCCAACTCTTTTGTTCTGAG 91374583

WT 91374584 CGAATGTGGAGACTGATTTTGGCAATGTTGAGAGCTGTGTTATCAGCAGTTTCACATTGACCCTTCCTCTGGTCAGCCTTCTTTAAACTTTGTAATCTGCCTTAGGACCCAGGCTTTTCTTTTTTTTTCTTTTTTCTTTCTTTCTTTCTTT 91374733
KI$^{cas9}$ 91374584 CGAATGTGGAGACTGATTTTGGCAATGTTGAGAGCTGTGTTATCAGCAGTTTCACATTGACCCTTCCTCTGGTCAGCCTTCTTTAAACTTTGTAATCTGCCTTAGGACCCAGGCTTTTCTTTTTTTTTCTTTTTTCTTTCTTTCTTTCTTT 91374733

WT 91374734 CTTTTTTTTTTTTTTTTTTTGTAGCAGCTTCTTTTGGTGATTGTCTTTTTTAAAAAGGGAAAGTACGTCTTTTTTTTTAAAACACTGCCATTAAGTCCATGGGACTCTTCAATTTTCACTTTCTATCAAGGGGAGTAATTTATTTGTGGT 91374883
KI$^{cas9}$ 91374734 CTTTTTTTTTTTTTTTTTTTGTAGCAGCTTCTTTTGGTGATTGTCTTTTTTAAAAAGGGAAAGTACGTCTTTTTTTTTAAAACACTGCCATTAAGTCCATGGGACTCTTCAATTTTCACTTTCTATCAAGGGGAGTAATTTATTTGTGGT 91374883

WT 91374884 TTTTGTGACATGACGTATTGAACATGGAAGTCAGGCTTCATAACAGTTACAGACCTTGTTCTGAGGATAGCTGGGATAATTCAGTACCAATTGATGTAAATTGTTCATTCTTCTGAGTATAGAAAGAGTCCCCAGAAATTATCTAAAAAT 91375033
KI$^{cas9}$ 91374884 TTTTGTGACATGACGTATTGAACATGGAAGTCAGGCTTCATAACAGTTACAGACCTTGTTCTGAGGATAGCTGGGATAATTCAGTACCAATTGATGTAAATTGTTCATTCTTCTGAGTATAGAAAGAGTCCCCAGAAATTATCTAAAAAT 91375033

WT 91375034 AGTAGTTTTATGTTTTTAAAAAAAATAAAGTTTTGCCTGTCTACTGCTGTAAGTTCTTCTGTTTCCTCCCGTGTCTGTGGTTTGTGCTGGGTTGGAGTGGGGAATCTCAGGAATTACTGCAGGCCAAGGGGGCTGACACAGTAAATGCTC 91375183
KI$^{cas9}$ 91375034 AGTAGTTTTATGTTTTTAAAAAAAATAAAGTTTTGCCTGTCTACTGCTGTAAGTTCTTCTGTTTCCTCCCGTGTCTGTGGTTTGTGCTGGGTTGGAGTGGGGAATCTCAGGAATTACTGCAGGCCAAGGGGGCTGACACAGTAAATGCTC 91375183

WT 91375184 AACTGAGCTTTGGTGATGCTCACTGCTGCGCGGTGATGTTGGCGACAGCTGCCTTTATATCATCCCCAGTATCCAGCTCGGGGCCAGGCACACAGGCTGCCCCTGCCTAGACAGAAGCTCAGTGAAGAATGGATGCCGCCAGCCAGCATG 91375333
KI$^{cas9}$ 91375184 AACTGAGCTTTGGTGATGCTCACTGCTGCGCGGTGATGTTGGCGACAGCTGCCTTTATATCATCCCCAGTATCCAGCTCGGGGCCAGGCACACAGGCTGCCCCTGCCTAGACAGAAGCTCAGTGAAGAATGGATGCCGCCAGCCAGCATG 91375333

WT 91375334 CGGGATTCCCTTCCACCCTGTAACACCAGAGCTCATTTCATGTCTTTTCTGAAATGCTCTGGGTATTTGACATTCTTTCGTCCTTCTTTCATATGGTTACCCTCAGAAGGAGCCACATGAATTTGCCTTTTTGCGCACGTGCCCGTGTTC 91375483
KI$^{cas9}$ 91375334 CGGGATTCCCTTCCACCCTGTAACACCAGAGCTCATTTCATGTCTTTTCTGAAATGCTCTGGGTATTTGACATTCTTTCGTCCTTCTTTCATATGGTTACCCTCAGAAGGAGCCACATGAATTTGCCTTTTTGCGCACGTGCCCGTGTTC 91375483

WT 91375484 TTTGTTTGGTGCCAGACAGGACACAAGTTTTGAACCTTTCGTAGTTAAAAGCTAATCAACCCAGCGTTCCTAGGCCAGCGCCTAGCACTGCAGCAGTGCTCAGGGCTCTTTTGGAGGGAGGAGGTACGATGGTTATTTCTTAGCCTAGCC 91375633
KI$^{cas9}$ 91375484 TTTGTTTGGTGCCAGACAGGACACAAGTTTTGAACCTTTCGTAGTTAAAAGCTAATCAACCCAGCGTTCCTAGGCCAGCGCCTAGCACTGCAGCAGTGCTCAGGGCTCTTTTGGAGGGAGGAGGTACGATGGTTATTTCTTAGCCTAGCC 91375633

WT 91375634 AGTTGGTGCCTGGTCGGAGTGATGAATGGTGCAGTGTGGGAAAACTAGCCTCCGCTGGGCTCCTCAGCAGTCATTTAGGGGTGGTTCTAGAGTTGTCTGAAACGGACTACACTCGGGGCTGCTGCTTGTGGTCTCGGGTATCTTGTCCAT 91375783
KI$^{cas9}$ 91375634 AGTTGGTGCCTGGTCGGAGTGATGAATGGTGCAGTGTGGGAAAACTAGCCTCCGCTGGGCTCCTCAGCAGTCATTTAGGGGTGGTTCTAGAGTTGTCTGAAACGGACTACACTCGGGGCTGCTGCTTGTGGTCTCGGGTATCTTGTCCAT 91375783

WT 91375784 GTCAAAGCTTGCCTTGTTTCTTTACTTGGAATCTGAAAAATGCTCAGGCTCAGACTCCTTCTCAAGTTGCATTTGGTCTCATCACCCTTGGCTTCCAGGGGCCTGAGACAATGGGACTGTTTATTGTCCTTGTCTTTTTTTACTCTTTTG 91375933
KI$^{cas9}$ 91375784 GTCAAAGCTTGCCTTGTTTCTTTACTTGGAATCTGAAAAATGCTCAGGCTCAGACTCCTTCTCAAGTTGCATTTGGTCTCATCACCCTTGGCTTCCAGGGGCCTGAGACAATGGGACTGTTTATTGTCCTTGTCTTTTTTTACTCTTTTG 91375933

WT 91375934 GAGTTCAACTAATTTAACCCACAATAAAGTTAATCCTCACTTCTTTGGGACAGTCAAGACTAACATATTAGCTGTTTTCAGGACCATTTTCATGAAAGCAAGTGTGTTGTTCCACTGGCTCTCGGGGCTGACATATGCAGTTGATGGCTG 91376083
KI$^{cas9}$ 91375934 GAGTTCAACTAATTTAACCCACAATAAAGTTAATCCTCACTTCTTTGGGACAGTCAAGACTAACATATTAGCTGTTTTCAGGACCATTTTCATGAAAGCAAGTGTGTTGTTCCACTGGCTCTCGGGGCTGACATATGCAGTTGATGGCTG 91376083

WT 91376084 TTTAGAAGTCAGGTTTCAGAGCAAACCACAACTGCCCTGGGTTACTGAAGTTCTCTTTAATGTTGGCTGTGAGCATGTGTGTGTCAAACATGAGGGAGTGGGATTACCTTACTATGTGTATGTTAGAGGGAGCAAGGGAGGGAGGGAGCA 91376233
KI$^{cas9}$ 91376084 TTTAGAAGTCAGGTTTCAGAGCAAACCACAACTGCCCTGGGTTACTGAAGTTCTCTTTAATGTTGGCTGTGAGCATGTGTGTGTCAAACATGAGGGAGTGGGATTACCTTACTATGTGTATGTTAGAGGGAGCAAGGGAGGGAGGGAGCA 91376233

WT 91376234 AAAGAAGGAGGGAAGTGTGTGTGTGTGTGTGTGTGTATGTGTGTGCACATGAGAGGGAGAGGGAGAGAGAGAGAACATGCTGTTGGAATTGAAGCTTTTACTGTCAATGAAAGCACACTGGTTAAATGCCTAGATGCCAGGGCGGGGGTACG 91376383
KI$^{cas9}$ 91376234 AAAGAAGGAGGGAAGTGTGTGTGTGTGTGTGTGTGTATGTGTGTGCACATGAGAGGGAGAGGGAGAGAGAGAGAACATGCTGTTGGAATTGAAGCTTTTACTGTCAATGAAAGCACACTGGTTAAATGCCTAGATGCCAGGGCGGGGGTACG 91376383

WT 91376384 TGGATCTGAGTACTGGCTTTGCTGCTTTCTAATTATGTAACTTAAGGCAAGTTGCTGAATTTCTCTGTTCCTTCCTTTTCCTGTCTTATCAAGTGAACAAAACTATGGTAGCGGCATTGAGGTTGCTCTAAGGCTCCAGTGTGTTGATTT 91376533
KI$^{cas9}$ 91376384 TGGATCTGAGTACTGGCTTTGCTGCTTTCTAATTATGTAACTTAAGGCAAGTTGCTGAATTTCTCTGTTCCTTCCTTTTCCTGTCTTATCAAGTGAACAAAACTATGGTAGCGGCATTGAGGTTGCTCTAAGGCTCCAGTGTGTTGATTT 91376533

WT 91376534 GGATAAACTGTTTAGAACAAAGCTTTGTGCGTAACCAGTCTTTACTCGCCATTTCTAGCAGCAATTAGCAATTGCACAAAATTAGAAGAAGAGGAAACATAGTCGTTAAGCCATGTCAAGCATGGCCATGCGTGTGAAATGCTTTAGGAG 91376683
KI$^{cas9}$ 91376534 GGATAAACTGTTTAGAACAAAGCTTTGTGCGTAACCAGTCTTTACTCGCCATTTCTAGCAGCAATTAGCAATTGCACAAAATTAGAAGAAGAGGAAACATAGTCGTTAAGCCATGTCAAGCATGGCCATGCGTGTGAAATGCTTTAGGAG 91376683

WT 91376684 CTATGGCTAGTCCTGCGGGTTTGTGTTACGAGTTTAGGGAGGTGTCTGTTGTAACAAAAACTACGTGAAGATTCTGCATGACGTCTTAAATGTAGTTTTGATTTCTTGCTTATGTAGTAGCGAGGGATACTTGGCTTCATGCATTCTGTC 91376833
KI$^{cas9}$ 91376684 CTATGGCTAGTCCTGCGGGTTTGTGTTACGAGTTTAGGGAGGTGTCTGTTGTAACAAAAACTACGTGAAGATTCTGCATGACGTCTTAAATGTAGTTTTGATTTCTTGCTTATGTAGTAGCGAGGGATACTTGGCTTCATGCATTCTGTC 91376833

WT 91376833 AGTAGGAACCTTGGCGGGCTCCACAGAAGGCTGCAGAGGTCACCTCAGGTTATCTCCGCTGAAGCCAGCGGGGAGGGAGAGCAGGCAGGGCCGTGTGTAGGAGCGTTTTTATGGGCCAGCTCTGGCTGTGGCACTCAACACATCTGTTCC 91376983
KI$^{cas9}$ 91376833 AGTAGGAACCTTGGCGGGCTCCACAGAAGGCTGCAGAGGTCACCTCAGGTTATCTCCGCTGAAGCCAGCGGGGAGGGAGAGCAGGCAGGGCCGTGTGTAGGAGCGTTTTTATGGGCCAGCTCTGGCTGTGGCACTCAACACATCTGTTCC 91376983

WT 91376983 CACTCCTCTGGCCAGAGCTGGGCGGGAAGGAGGAGAGCGAGCAGGAGCTGCCGTTCGTCCTCCGCATGTTCCTCAGGAGTCTAGAGATGGAGAAGTGAGAAAAGCCTTGTCTCCAGACTCTGAACTGAAACTGCACAACTGTCCCATTGG 91377133
KI$^{cas9}$ 91376983 CACTCCTCTGGCCAGAGCTGGGCGGGAAGGAGGAGAGCGAGCAGGAGCTGCCGTTCGTCCTCCGCATGTTCCTCAGGAGTCTAGAGATGGAGAAGTGAGAAAAGCCTTGTCTCCAGACTCTGAACTGAAACTGCACAACTGTCCCATTGG 91377133

WT 91377134 GGGTTTGTGAAATGCAGTCTTACAGCAGCTTCCAAAGATGACACTTATTTTATGCACACACACACAAAAAAGTGACTACATTGGATAATAATTTGTAAGTGTTTTCTAATTTCCAAGAGAGGTTGCTCTAGGGCTCCAGAGTGTTGATTT 91377283
KI$^{cas9}$ 91377134 GGGTTTGTGAAATGCAGTCTTACAGCAGCTTCCAAAGATGACACTTATTTTATGCACACACACACAAAAAAGTGACTACATTGGATAATAATTTGTAAGTGTTTTCTAATTTCCAAGAGAGGTTGCTCTAGGGCTCCAGAGTGTTGATTT 91377283

WT 91377284 GGAGTTTCTAAGAAAATCATTAAGGGGTTTTCCAGGCCAAATAAGTTTTAAAGTAAAATGTTTTAAAAAACTGCTCACTTTCTTAAATTGTAAAGAAGTGTTAATGGGAAATGTGGGCGAAATTTAATCTGTTTATCCTGCTGGTGAGTG 91377433
KI$^{cas9}$ 91377284 GGAGTTTCTAAGAAAATCATTAAGGGGTTTTCCAGGCCAAATAAGTTTTAAAGTAAAATGTTTTAAAAAACTGCTCACTTTCTTAAATTGTAAAGAAGTGTTAATGGGAAATGTGGGCGAAATTTAATCTGTTTATCCTGCTGGTGAGTG 91377433

WT 91377434 GGCCTTCAAAGGCCTCAGGGTGTGGTTTGCCAACAAAGGCCTTCAGCGGCCCTGCTGGTGGCTCGCGGTCACATTTTGCATTCTGAGCCTTTAGATATTTTGCTCCCTTACATTCCTCTCTTTTATTCCTTTATTTTCTGATAAAATATT 91377583
KI$^{cas9}$ 91377434 GGCCTTCAAAGGCCTCAGGGTGTGGTTTGCCAACAAAGGCCTTCAGCGGCCCTGCTGGTGGCTCGCGGTCACATTTTGCATTCTGAGCCTTTAGATATTTTGCTCCCTTACATTCCTCTCTTTTATTCCTTTATTTTCTGATAAAATATT 91377583

WT 91377584 TGCTCCGGTGAAACCTAAGAACAGGAATCACACTGCATTTAATAACACAGGGGCTTGGTCTGGTGAATTCCGTGCCAGAGAGGAACAGTGTCTTGTGTGGTTTTGTC 91377690
KI$^{cas9}$ 91377584 TGCTCCGGTGAAACCTAAGAACAGGAATCACACTGCATTTAATAACACAGGGGCTTGGTCTGGTGAATTCCGTGCCAGAGAGGAACAGTGTCTTGTGTGGTTTTGTC 91377690
