## Supplementary material for "The rs1421085 variant within *FTO* promotes but not inhibits thermogenesis and is potentially associated with human migration": table S1-S6

**Table S1. Off-target Sites Prediction Top 10**

| **Coordinates** | **strand** | **MM*** | **target_sequence** | **PAM** | **distance** |  | **gene name** |
| --- | --- | --- | --- | --- | --- | --- | --- |
| chr8:91374360-91374382 | - | 0 | CTTAATCA[ATACGATGCCTT] | AGG | 13354 | I | Fto |
| chr3:5498973-5498995 | - | 4 | GTTTATAA[ATATGATGCCTT] | AGG | 14000 | - | 4930555M17Rik |
| chr14:39161796-39161818 | - | 4 | CTAAAACA[TGACGATGCCTT] | TGG | 58253 | I | Gm20642 |
| chr10:50467540-50467562 | - | 4 | ATGAATCC[ATAAGATGCCTT] | TGG | NA | - | NA |
| chr17:93375023-93375045 | + | 4 | CTGAATAA[TGACGATGCCTT] | CGG | NA | - | NA |
| chr11:90012399-90012421 | - | 4 | CTCCATCT[ATATGATGCCTT] | GGG | 9505 | - | Pctp |
| chr7:91069018-91069040 | + | 4 | ATTAAACC[ATATGATGCCTT] | AGG | 21666 | - | Dlg2 |
| chr2:117820099-117820121 | - | 4 | CTTGAGTA[ATATGATGCCTT] | GGG | 1807 | - | 4930412B13Rik |
| chr15:82106342-82106364 | - | 4 | CTCAAACA[AACCGATGCCTT] | TGG | 666 | I | Mei1 |
| chr10:130452922-130452944 | + | 4 | CTAAACAA[ATACCATGCCTT] | TGG | 0 | E | Vmn2r86 |

* MM: Mouse Musculus GRCm38/mm10

Data Analyze: <https://cctop.cos.uni-heidelberg.de:8043/>

Referrence: https://cctop.cos.uni-heidelberg.de:8043/help.html

**Table S2. Primers**

| **Genotyping Primers** | **sequence (5'→3'）** |
| --- | --- |
| mouse Fto-F | AGCCCAGCAAACTCATTCCT |
| mouse Fto-R | CAGATTAAGGTGACGGGCTGGAT |
| Fto sequencing | AAGGTGACATACACCAGGAGCC |
| loxP-F | GCCGATTAGCAGAGAAGTGATCAG |
| loxP-R | CTTGAACATCTTCCTCCACCTGAG |
| Cre-F | ATTTGCCTGCATTACCGGTCG |
| Cre-R | CAGCATTGCTGTCACTTGGTC |
| **qPCR Primers** | **sequence (5'→3'）** |
| 36B4-F | GAAACTGCTGCCTCACATCCG |
| 36B4-R | GCTGGCACAGTGACCTCACACG |
| Ucp1-F | AGGCTTCCAGTACCATTAGGT |
| Ucp1-R | CTGAGTGAGGCAAAGCTGATTT |
| Fto-F | GACACTTGGCTTCCTTACCTG |
| Fto-R | CTCACCACGTCCCGAAACAA |
| Cox8b-F | GAACCATGAAGCCAACGACT |
| Cox8b-R | GCGAAGTTCACAGTGGTTCC |
| Cox7a-F | CAGCGTCATGGTCAGTCTGT |
| Cox7a-R | AGAAAACCGTGTGGCAGAGA |
| Elovl6 F | CCCGAACTAGGTGACACGAT |
| Elovl6 R | CCAGCGACCATGTCTTTGTA |
| C/EBPα F | GGTTTCGGGTCGCTGGATCTCTAG |
| C/EBPα-R | ACGGCCTGACTCCCTCATCTTAGAC |
| C/ebpβ-F | TTATAAACCTCCCGCTCGGC |
| C/ebpβ-R | TTCCATGGGTCTAAAGGCGG |
| Pgc1α-F | AGCCGTGACCACTGACAACGAG |
| Pgc1α-R | RGCTGCATGGTTCTGAGTGCTAAG |
| Cidea-F | TGCTCTTCTGTATCGCCCAGT |
| Cidea-R | GCCGTGTTAAGGAATCTGCTG |
| Pparγ2-F | GCATGGTGCCTTCGCTGA |
| Pparγ2-R | TGGCATCTCTGTGTCAACCATG |
| Dio2-F | AATTATGCCTCGGAGAAGACCG |
| Dio2-F | GGCAGTTGCCTAGTGAAAGGT |
| Prdm16-F | TGACGGATACAGAGGTGTCAT |
| Prdm16-R | ACGCTACACGGATGTACTTGA |
| Irx3-F | GGCAATGCTTATGGGAGCGA |
| Irx3-R | CGCTGTCTAAGTTTTCCAAATCG |
| Irx5-F | GCACGGATGAGCTCGGCCGCTC |
| Irx5-R | GGGTGATATCCCAAGGAACCTG |
| Irx6-F | CTCAGTATGAGTTCAAGGATGCTG |
| Irx6-R | CTCCCTTGTGGCATTCTTCCTG |
| Fabp4-F | ACACCGAGATTTCCTTCAAACTG |
| Fabp4-R | CCATCTAGGGTTATGATGCTC |
| Rpgrip1l-F | ACTGGAAGACAGATTTTTGCGT |
| Rpgrip1l-R | AACTAGCCGTATTAACTTGGTGG |

**Table S3. Primer Sequences**

| **Lentiviruse siRNA** | | | | |
| --- | --- | --- | --- | --- |
| **Marker** | **Gene** | **Gene ID** | **TargetSequence** | **GC%** |
| shFTO-1 | Fto | NM_011936 | GCTTGAAGACACTTGGCTT |  |
| shFTO-2 | Fto | NM_011936 | GCATGTCAGACCTTCCTAA |  |
| shFTO-3 | Fto | NM_011936 | GCTGAGGCAGTTCTGGTTT |  |
| shCtrl | NC |  | TTCTCCGAACGTGTCACGT | 52.6 |
| **shRNA and Primers** | | | | |
| **DNA primers** | **Sequence** | | | |
| shFTO-1 F | CCGGGCTTGAAGACACTTGGCTTCTCAAGAGAAAGCCAAGTGTCTTCAAGCTTTTTTG | | | |
| shFTO-1 R | AATTCAAAAAAGCTTGAAGACACTTGGCTTTCTCTTGAGAAGCCAAGTGTCTTCAAGC | | | |
| shFTO-2 F | CCGGGCATGTCAGACCTTCCTAATTCAAGAGATTAGGAAGGTCTGACATGCTTTTTTG | | | |
| shFTO-2 R | AATTCAAAAAAGCATGTCAGACCTTCCTAATCTCTTGAATTAGGAAGGTCTGACATGC | | | |
| shFTO-3 F | CCGGGCTGAGGCAGTTCTGGTTTCTCAAGAGAAAACCAGAACTGCCTCAGCTTTTTTG | | | |
| shFTO-3 R | AATTCAAAAAAGCTGAGGCAGTTCTGGTTTTCTCTTGAGAAACCAGAACTGCCTCAGC | | | |
| shCtrl F | CCGGTTCTCCGAACGTGTCACGTTTCAAGAGAACGTGACACGTTCGGAGAATTTTTTG | | | |
| shCtrl R | AATTCAAAAAATTCTCCGAACGTGTCACGTTCTCTTGAAACGTGACACGTTCGGAGAA | | | |
| CMV-F | CGCAAATGGGCGGTAGGCGTG | | | |

**Table S4. Study details**

| **FTO & BMI> 8** | | | | | | | |
| --- | --- | --- | --- | --- | --- | --- | --- |
| **Number** | **Auther** | **Cohort name** | **Publication year** | **Country** | **Ethnicity** | **Genotype** | **DOI** |
| 1 | Josefin A. Jacobsson | National Childhood Obesity Centre at Karolinska University Hospital, | 2008 | Sweden | Swedish | rs9939609 | DOI: 10.1016/j.bbrc.2008.01.087 |
| 2 | I.Labayen | The HELENA cross- sectional study | 2016 | Vitoria, Spain | European | rs9939609 | DOI: 10.1016/j.numecd.2016.07.010 |
| 3 | Liliane dos SantosRodrigues | RPS Cohort of São Luís, Maranhão. | 2019 | Brazil | Brazilian | rs9939609 | DOI: 10.1016/j.jped.2019.05.006 |
| 4 | Gaifen Liu | Georgia Cardiovascular Twin study the LACHY study and the APEX study | 2010 | The Netherlands | European whites and African American | rs9939609 | DOI: 10.1186/1471-2350-11-57 |
| 5 | Bo Xi | Beijing Child and Adolescent Metabolic Syn- drome (BCAMS) study | 2010 | China | Chinese | rs9939609 | DOI: 10.1186/1471-2350-11-107 |
| 6 | Hongyun Fang | NA^a^ | 2010 | China | Chinese | rs9939609 | DOI: 10.1186/1471-2350-11-136 |
| 7 | Josune Olza | NA | 2013 | Spain | European descent | rs9939609 | DOI: 10.1186/1471-2350-14-123 |
| 8 | Marian Tanofsky-Kraff | NA | 2009 | US | Non-Hispanic white and others | rs9939609 | DOI: 10.3945/ajcn.2009.28439 |
| 9 | Lizbeth González-Herrera | NA | 2019 | México | Mayan | rs1421085 | DOI: 10.1002/ajhb.23192 |
| 10 | S K Vasan | NA | 2013 | South India | Hiefly Dravidian in origin, with 1.2% being a mixture of Punjabi and Marwari ethnicity | rs9939609 | DOI: 10.1111/j.2047-6310.2013.00118.x |
| 11 | Maarit Hakanen | STRIP study | 2009 | Finland | Finn | rs9939609 | DOI: 10.1210/jc.2008-1199 |
| 12 | Femke Rutters | Dutch Caucasian cohort | 2011 | Dutch | Dutch Caucasian | rs9939609 | DOI: 10.1210/jc.2010-2413 |
| 13 | David Albuquerque | NA | 2013 | Portuguese | Portuguese | rs9939609 | DOI: 10.1371/journal.pone.0054370 |
| 14 | Junqing Wu | NA | 2014 | China | Chinese | rs9939609 | DOI: 10.1371/journal.pone.0098984 |
| 15 | Min Yang | NA | 2014 | China | Chinese | rs9939609 | DOI: 10.1371/journal.pone.0104574 |
| 16 | Robert A Scott | Growth, Exercise and Nutrition Epidemiological Study in preSchoolers (GENESIS) | 2010 | Greece | Greek | rs17817449 | DOI: 10.1038/ejhg.2010.131 |
| 17 | Zdenka Pausova | Saguenay Youth Study | 2009 | United Kingdom | Saguenay-Lac St Jean(French Canadian Founder Population) | rs9939609 | DOI: 10.1161/CIRCGENETICS.109.857359 |
| 18 | Ana Maria Obregón Rivas | NA | 2018 | Chile | Chilean | rs9939609 | DOI: 10.1016/j.nut.2018.03.001 |
| 19 | Keiko Shinozaki | Shunan Child Health Cohort Study | 2018 | Japan | Japanese | rs1558902 | DOI: 10.1111/ped.13578 |
| 20 | Jane Wardle | Twins’ Early Development Study（TEDS ） | 2008 | UK | United Kingdom children | rs9939609 | DOI: 10.1210/jc.2008-0472 |
| 21 | Harald Mangge | STYrian Juvenile OBesity Study (STYJOBS) | 2011 | Australia | European | rs9939609 | DOI: 10.1155/2011/186368 |
| ^a^. Not available. | | | | | | | |
| **FTO & BMI ≤ 8** | | | | | | | |
| **Number** | **Auther** | **Cohort name** | **Publication year** | **Country** | **Ethnicity** | **Genotype** | **DOI** |
| 1 | D O Mook-Kanamori | he Generation R Study | 2011 | Dutch | Dutch ethnicity | rs9939609 | DOI: 10.1007/BF03346689 |
| 2 | Meixian Zhang | Beijing Child and Adolescent Metabolic Syndrome (BCAMS) study | 2014 | China | Chinese | rs9939609 | DOI: 10.1371/journal.pone.0097545 |
| 3 | Fabio Lauria | IDEFICS | 2012 | Italy | White European descent | rs9939609 | DOI: 10.1371/journal.pone.0048876 |
| 4 | Timothy M. Frayling | The Avon Longitudinal Study of Parents and Children (ALSPAC) cohort and the Northern Finland 1966 birth cohort (NFBC1966) | 2008 | England and Finland | English and Finn | rs9939609 | DOI: 10.1126/science.1141634 |
| **FTO& Birthweigt** | | | | | | | |
| **Number** | **Auther** | **Cohort name** | **Publication year** | **Country** | **Ethnicity** | **Genotype** | **DOI** |
| 1 | Abel López-Bermejo | Neonatal Unit of the Obstetrics and Gynecology Department of the Hospital Sant Joan de De ́u, | 2008 | Spain | Spanish | rs9939609 | DOI: 10.1210/jc.2007-2343 |
| 2 | Maarit Hakanen | STRIP study | 2009 | Finland | Finn | rs9939609 | DOI: 10.1210/jc.2008-1199 |
| 3 | Bo Xi | Beijing Child and Adolescent Metabolic Syn- drome (BCAMS) study | 2010 | China | Chinese | rs9939609 | DOI: 10.1186/1471-2350-11-107 |
| 4 | Nuananong Seal | NA | 2011 | America | American Indian | rs9939609 | DOI: 10.1089/gtmb.2010.0188 |
| 5 | Tanaka | NA | 2012 | Japan | Japanese | rs1558902 | DOI: 10.5551/jat.11940 |
| 6 | Fleur P Velders | Generation R Study | 2012 | The Netherlands | Northern European descent | rs9939609 | DOI: 10.1371/journal.pone.0049131 |
| 7 | Olivier S Descamps | NA | 2014 | Belgium | European whites | rs9939609 | DOI: 10.1186/s12863-014-0145-0 |
| 8 | Elina Molou | NA | 2015 | Greece | Greek | rs9939609 | DOI: 10.1515/jpem-2014-0320 |
| 9 | Eva Gesteiro | NA | 2016 | Spain | Caucasian | rs9939609 | DOI: 10.1007/s13105-016-0467-7 |
| 10 | Claudiu Mărginean | Obstetrics Gynecology Tertiary Hospital from Romania | 2016 | Romania | Romanian | rs9939609 | DOI: 10.1097/MD.0000000000005551 |
| 11 |  | GOCY^b^ | 2020 | China | Chinese | rs1421085 |  |
| ^b^. data from the Genetics of Obesity in Chinese Youngs (GOCY) study. | | | | | | | |

**Table S5. Study details**

| **Populations** | **Study Fullname** | **Full name** |
| --- | --- | --- |
| JPT | 1000Genomes | Japanese in Tokyo, Japan |
| KR | KOREAN population from KRGDB | KOREAN |
| CHB | 1000Genomes | Han Chinese in Beijing, China |
| CHS | 1000Genomes | Southern Han Chinese |
| KHV | 1000Genomes | Kinh in Ho Chi Minh City, Vietnam |
| VI | Vietnamese | Vietnamese |
| CDX | 1000Genomes | Chinese Dai in Xishuangbanna, China |
| BEB | 1000Genomes | Bengali from Bangladesh |
| SI | Siberian | Siberian |
| PJL | 1000Genomes | Punjabi from Lahore, Pakistan |
| QAT | Qatari | Qatari |
| DAG | Genome-wide autozygosity in Daghestan | Daghestan |
| LWK | 1000Genomes | Luhya in Webuye, Kenya |
| FIN | 1000Genomes | Finnish in Finland |
| EST | Genetic variation in the Estonian population | Estonian |
| NS | Northern Sweden | Northern Sweden |
| GWD | 1000Genomes | Gambian in Western Divisions in the Gambia |
| DA | The Danish reference pan genome | Danish |
| TSI | 1000Genomes | Toscani in Italia |
| ESN | 1000Genomes | Esan in Nigeria |
| NL | Genome of the Netherlands Release 5 | Netherlands |
| YRI | 1000Genomes | Yoruba in Ibadan, Nigeria |
| ITU | 1000Genomes | Indian Telugu from the UK |
| STU | 1000Genomes | Sri Lankan Tamil from the UK |
| GBR | 1000Genomes | British in England and Scotland |
| IBS | 1000Genomes | Iberian Population in Spain |
| MSL | 1000Genomes | Mende in Sierra Leone |
| ACB | 1000Genomes | African Caribbeans in Barbados |
| PUR | 1000Genomes | Puerto Ricans from Puerto Rico |
| CLM | 1000Genomes | Colombians from Medellin, Colombia |
| PEL | 1000Genomes | Peruvians from Lima, Peru |
| DOM | The PAGE Study | Dominican |
| CUB | The PAGE Study | Cuban |
| GIH | 1000Genomes | Gujarati Indian from Houston, Texas |
| ASW | 1000Genomes | Americans of African Ancestry in SW USA |
| MX | The PAGE Study | Mexican |
| CEU | 1000Genomes | Utah Residents (CEPH) with Northern and Western European Ancestry |
| MXL | 1000Genomes | Mexican Ancestry from Los Angeles USA |
| NHPI | The PAGE Study | Native Hawaiian |

**Table S6. Acronym**

| **Abbreviation** | **Full name** |
| --- | --- |
| AGE | agarose gel electrophoresis |
| ARID5b | AT-rich interactive domain 5B |
| ATCC | American type culture collection |
| BAT | brown adipose tissue |
| BMI | body mass index |
| BMR | basal metabolic rate |
| C/ebpα | CCAAT-enhancer binding protein α |
| Cidea | cell death-inducing DNA fragmentation factor alpha-like effector A |
| Cox7a | cytochrome c oxidase subunit 7A1 |
| Cox8b | cytochrome c oxidase subunit 8B |
| CUX1 | cut like homeobox 1 |
| Dio2 | typeⅡiodothyronine deiodinase |
| DMEM | Dulbecco's modified eagle medium |
| DMSO | dimethyl sulphoxide |
| EGG Consortium | Early Growth Genetics Consortium |
| Elovl6 | elongase of very long chain fatty acids family member 6 |
| ENT | entacapone |
| ES cell | embryonic stem cell |
| eWAT | epididymal white adipose tissues |
| F12 | Ham's F12 nutrient medium |
| FBS | fetal bovine serum |
| FTO/Fto | fat mass and obesity associated |
| GWAS | genome-wide association studies |
| H&E | hematoxylin and eosin stain |
| HDL-c | high-density lipoprotein-cholesterol |
| HFD | high fat diet |
| iBAT | interscapular BAT |
| IBMX | 3-isobutyl-1-methylxanthine |
| IPGTT | intraperitoneal glucose-tolerance tests |
| IRX3/Irx3 | iroquois homeobox 3 |
| ITT | insulin tolerance test |
| iWAT | inguinal white adipose tissue |
| KI | knock-in |
| LDL-c | low-density lipoprotein-cholesterol |
| LGM | Last Glacial Maximum |
| m6A | N6-methyladenosine |
| NCD | normal chow diet |
| OCR | oxygen consumption rate |
| PBS | phosphate buffer solution |
| PCR | polymerase chain reaction |
| Pgc-1α | peroxisome proliferator activated receptor coactivator-1 alpha |
| PLIN1 | perilipin1 |
| Prdm16 | PR domain containing 16 |
| RER | respiratory exchange ratio |
| RIP | RNA immunoprecipitation |
| Rosi | rosiglitazone |
| SNP | single nucleotide polymorphism |
| SVF | stromal vascular fraction |
| T3 | triiodothyronine |
| TC | total cholesterol |
| TEM | transmission electron microscopy |
| TG | triglyceride |
| UCP1 | uncoupling protein 1 |
| WT | wild-type |
| △iBAT | excision of interscapular BAT |
